# β-arrestin-independent endosomal cAMP signaling by a polypeptide hormone GPCR

**DOI:** 10.1101/2022.09.07.506997

**Authors:** Emily E. Blythe, Mark von Zastrow

## Abstract

Many GPCRs are now recognized to initiate a second phase of G protein (G_s_) -dependent signaling through the cAMP cascade after endocytosis. A prevailing current view is that endocytosis-promoted signaling from GPCRs is inherently β-arrestin-dependent because β-arrestin is necessary for receptors to internalize and, for some GPCRs, it also promotes G_s_ activation on endosomes. Here we revise this view by showing that the vasoactive intestinal peptide receptor 1 (VIPR1), a prototypic secretin-family polypeptide hormone receptor, remains bound to β-arrestin after endocytosis but does not require β-arrestin either to internalize or to generate an endosomal signal. β-arrestin instead resolves the endosomal signal into a temporally separated cAMP peak, and it does so by attenuating signaling from the plasma membrane without detectably affecting the endosomal response. The mechanistic basis for this location-specific difference in β-arrestin function is the formation of distinct VIPR1/β-arrestin complexes at each location. The signal-attenuating complex formed at the plasma membrane does not require GRK-mediated phosphorylation of receptors, while the signaling-neutral complex present on the endosome membrane, in contrast, is GRK-dependent. To our knowledge, the present results provide the first direct demonstration that endosomal GPCR signaling can occur in the complete absence of β-arrestin. They also reveal a discrete role of β-arrestin in sculpting the spatiotemporal profile of cellular GPCR - G protein signaling through the location-specific formation or remodeling of GPCR/β-arrestin complexes.

## Introduction

G protein-coupled receptors (GPCRs) comprise the largest class of signaling receptors, regulate essentially every physiological process and are important drug targets^1^. Upon binding agonist ligands, GPCRs engage cognate heterotrimeric G proteins to transduce signaling through specific downstream effectors^2,3^. The biochemical basis of GPCR activation has been studied to a level of atomic detail^4,5^, but we are only now beginning to understand the subcellular organization of GPCR signaling^2,6,7^. Many GPCRs are not restricted to the plasma membrane and transit the endocytic pathway^8^. However, endocytosis was long believed only to impact the longer-term homeostatic regulation of GPCRs and not affect the response to acute agonist application. This view has changed due to the accumulation of substantial evidence that various GPCRs have the capacity to engage G proteins after endocytosis, as well as from the plasma membrane, and can leverage the endocytic network to promote or sustain cellular signaling^2,6,7^.

Support for this still-evolving view is well-developed for GPCRs that signal by coupling to stimulatory heterotrimeric G proteins (G_s_) which activate adenylyl cyclases to produce cAMP. A number of such GPCRs have now been shown to engage G_s_ on the endosome limiting membrane as well as the plasma membrane, enabling receptors to initiate sequential ‘waves’ of signaling from each location^2,6,9–15^. Numerous studies have demonstrated significant differences in the downstream effects of cAMP generated at the plasma membrane compared to internal membrane compartments, both at the cell and tissue levels^9,11,13–19^, highlighting how spatiotemporal aspects of GPCR activation can profoundly influence functional responses through cAMP. Yet, how such signaling diversity is programmed remains poorly understood.

GPCR-elicited cellular cAMP signaling can be determined by many factors, including the ligand’s binding affinity for receptors^10,11^ and specific features of the receptor’s trafficking itinerary^20,21^. One important factor is the interaction between GPCRs and β-arrestin. β-arrestins (β-arrestin-1 and β-arrestin-2) were discovered as scaffolding proteins which are recruited to and functionally desensitize activated receptors at the plasma membrane^22^. β-arrestins additionally function as essential endocytic adaptor proteins for many GPCRs, promoting receptor endocytosis via clathrin-coated pits and driving receptor delivery to endosomes^22,23^. Polypeptide hormone receptors provide particularly clear examples of such behavior. In particular, the thyroid stimulating hormone receptor (TSHR), parathyroid hormone receptor-1 (PTHR1) and vasopressin-2 receptor (V2R) have all been extensively shown to produce an endosomal cAMP signal that is inherently β-arrestin-dependent because β-arrestin is required for receptors to internalize^9–11,17,18,24–27^. In addition, for the PTHR1 and V2R, persistent binding to β-arrestin after internalization has been shown to sustain the endosomal cAMP signal^2,11,24–29^. Accordingly, a prevailing current view in the field is that GPCR signaling from endosomes is strictly β-arrestin-dependent. There has been evidence for many years that some GPCRs can internalize independently of β-arrestin^23,30^, and some of these have been shown to produce an endosomal cAMP signal (e.g.,^14^). However, it is not known if endosomal cAMP signaling from such GPCRs can actually occur in the absence of β-arrestin. Indeed, to our knowledge, it has not been determined if any GPCR can produce an endosomal G_s_-cAMP signal in the absence of β-arrestin. If so, a fundamental next question that arises is whether or how β-arrestin impacts the GPCR-elicited cellular cAMP response in such a case.

Here, we address these questions by focusing on the vasoactive intestinal peptide receptor 1 (VIPR1 or VPAC1) as a model secretin-subfamily polypeptide hormone GPCR shown previously to associate with β-arrestin both at the plasma membrane and endosome, but whose internalization is insensitive to a dominant negative mutant of β-arrestin^30,31^. We show that agonist-induced VIPR1 endocytosis is indeed β-arrestin-independent and then leverage this property to disentangle effects of β-arrestin binding and endocytosis on the cellular cAMP signal produced by endogenous receptor activation. While β-arrestin is clearly not required for endosomal G_s_ activation or cAMP signaling by VIPR1, we show that it plays a different role in temporally resolving the effects of VIPR1-G_s_ activation from the plasma membrane and endosomes into separate and sequential peaks of global cytoplasmic cAMP elevation. The present results provide the first direct example of a GPCR for which G_s_-cAMP signaling is fully β-arrestin-independent and reveal a discrete function of β-arrestin in spatiotemporally sculpting the cell’s overall cAMP response.

## Results

### VIPR1 is internalized in β-arrestin double knockout cells despite agonist-induced β-arrestin recruitment to both the plasma membrane and endosomes

To determine whether VIPR1 could serve as a model receptor to disentangle the effects of β-arrestin on trafficking and signaling, we first set out to characterize VIPR1 internalization in HEK293 cells. To do so, we used a cell-impermeant HaloTag dye (JF_635_i-HTL) to selectively label receptors in the plasma membrane of living cells^32^. As expected^30,31,33,34^, HaloTag-VIPR1 underwent endocytosis in wild type cells upon addition of its agonist, vasoactive intestinal peptide (VIP) (Fig. 1a). Surface-labeled receptors were visible in intracellular puncta that colocalized with both an overexpressed DsRed-labeled EEA1 and endogenous EEA1 (Extended Data Fig. 1a-c), identifying these structures as early endosomes. Some receptor-positive endosomes were visible even before VIP application, suggesting a low level of constitutive (agonist-independent) endocytosis. However, VIP markedly increased receptor localization in DsRed2-EEA1-positive endosomes, confirming that VIPR1 endocytosis is stimulated by agonist (Extended Data Fig. 1a). Agonist-induced internalization was further verified and quantified using flow cytometry, which showed an extremely rapid loss of surface receptor labeling that reached completion within 10 minutes of treatment with VIP (Fig. 1b). Both constitutive and agonist-induced components of VIPR1 internalization were observed under conditions in which VIPR1 expression level varies, though higher VIPR1 expression led to lower apparent levels of internalization (Extended Data Fig. 1d,e)^34^.

**Fig. 1.**
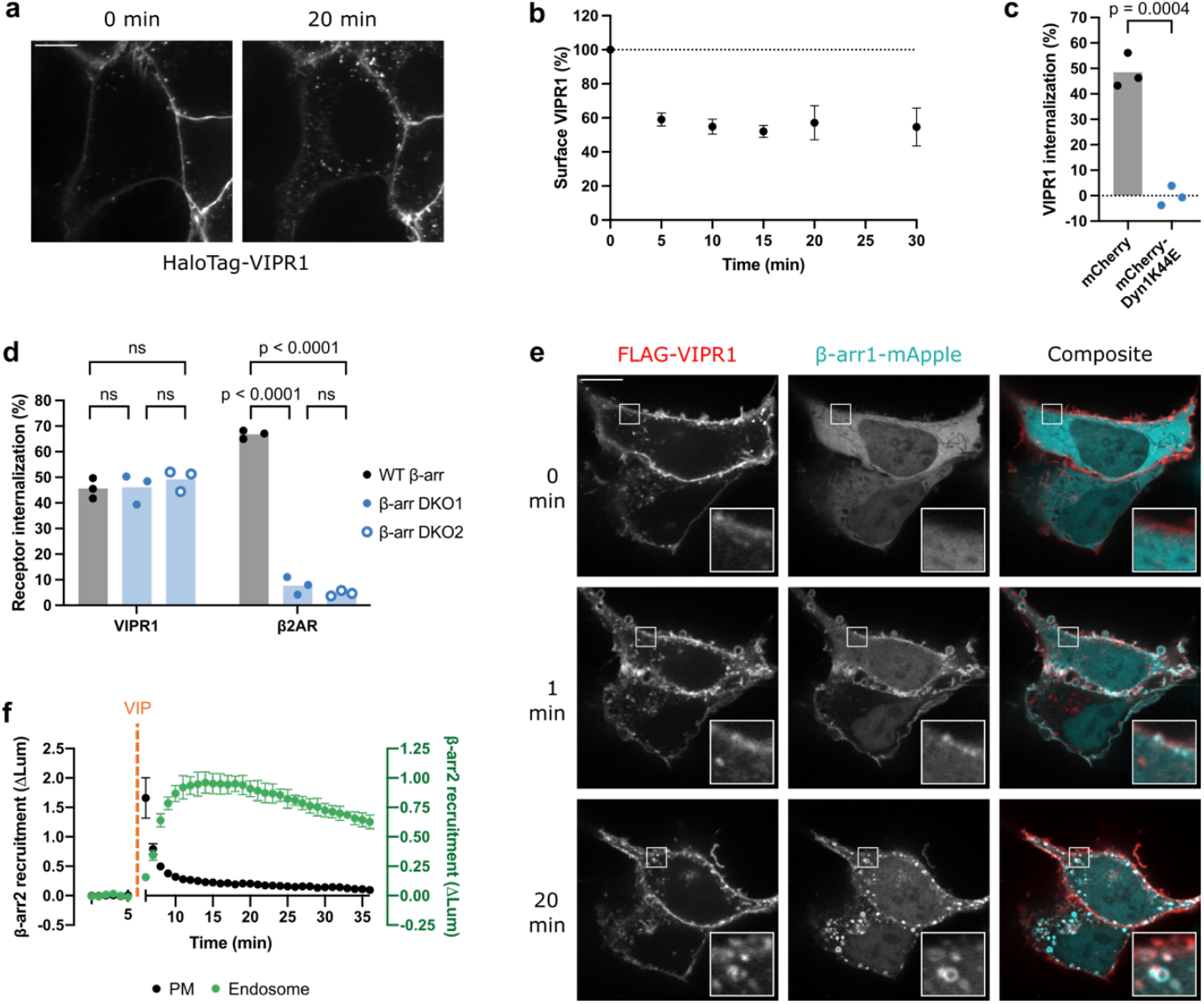
β-arrestin is recruited to activated VIPR1 but is not required for its internalization. **a**, Representative stills from time-lapse microscopy of cells expressing HaloTag-VIPR1 immediately before or 20 minutes after stimulation with 500 nM VIP. HaloTag-VIPR1 was expressed under a doxycycline(Dox)-inducible promoter, and surface receptor was labeled with cell-impermeant JF_635_i-HTL for 10 minutes prior to imaging. Scale bar is 10 μm. **b**, Time course of surface levels of HaloTag-VIPR1 after 500 nM VIP treatment, as measured by flow cytometry. **c**, Internalization of HaloTag-VIPR1 in cells co-expressing mCherry-Dyn1K44E or mCherry after a 30 minute treatment with 500 nM VIP, as measured by flow cytometry. Significance was determined by an unpaired t-test. **d**, Internalization of HaloTag-VIPR1 and HaloTag-β2AR after a 30 minute treatment with 500 nM VIP or a 20 minute treatment 1 μM isoproterenol, respectively, in cell lines expressing endogenous β-arrestin (WT β-arr) or lacking β-arrestin (β-arr DKO1 and β-arr DKO2). Receptor internalization was measured by flow cytometry, and significance was determined by a two-way ANOVA with Tukey’s multiple comparisons test. **e**, Representative stills from time-lapse microscopy of WT HEK293 cells co-expressing FLAG-VIPR1 and β-arr1-mApple, with 500 nM VIP added at 0 minutes. Surface FLAG-VIPR1 was labeled with anti-FLAG M1 antibody conjugated to Alexa Fluor 647 for 10 minutes before imaging. Scale bar is 10 μm. **f**, NanoBiT bystander assays showing recruitment of β-arr2-smBiT to the plasma membrane (LgBiT-CAAX, black) or the endosome (endofin-LgBiT, green), as measured by a change in luminescence upon addition of 1 µM VIP at 5 minutes. All data represent three biological replicates, shown as individual data points or mean ± s.d.

After establishing VIPR1 undergoes agonist-induced internalization, we next explored its basic mechanistic properties. Like many other GPCRs, and consistent with previous studies^30,31^, VIPR1 endocytosis was dynamin-dependent, as overexpression of a dominant negative mutant of dynamin 1 (Dyn1-K44E) fully suppressed agonist-induced internalization (Fig. 1c). Previous evidence relevant to β-arrestin dependence was based on insensitivity of VIPR1 to endocytic inhibition by dominant negative mutant constructs overexpressed on a wild type background^30,31^. In order to test the requirement for β-arrestin in VIPR1 internalization more incisively, we examined the effect of depleting endogenous β-arrestin. We generated two independent mutant HEK293 cell lines lacking both β-arrestin-1 (β-arr1) and β-arrestin-2 (β-arr2) using CRISPR. We validated complete β-arrestin knockout in both β-arrestin double knockout (β-arr DKO) cell lines biochemically, by genomic DNA sequencing and western blotting (Extended Data Fig. 2a,b), and functionally by loss of β2-adrenergic receptor (β2AR) internalization (a β-arrestin-dependent GPCR) (Fig. 2d). Robust internalization of VIPR1 was still observed in β-arr DKO cells, and complete depletion of β-arrestin did not affect the degree or kinetics of agonist-induced internalization in either independent cell line (Fig. 1d and Extended Data Fig. 2c). Therefore, agonist-induced endocytosis of VIPR1 proceeds through a dynamin-dependent, β-arrestin-independent mechanism.

**Fig. 2.**
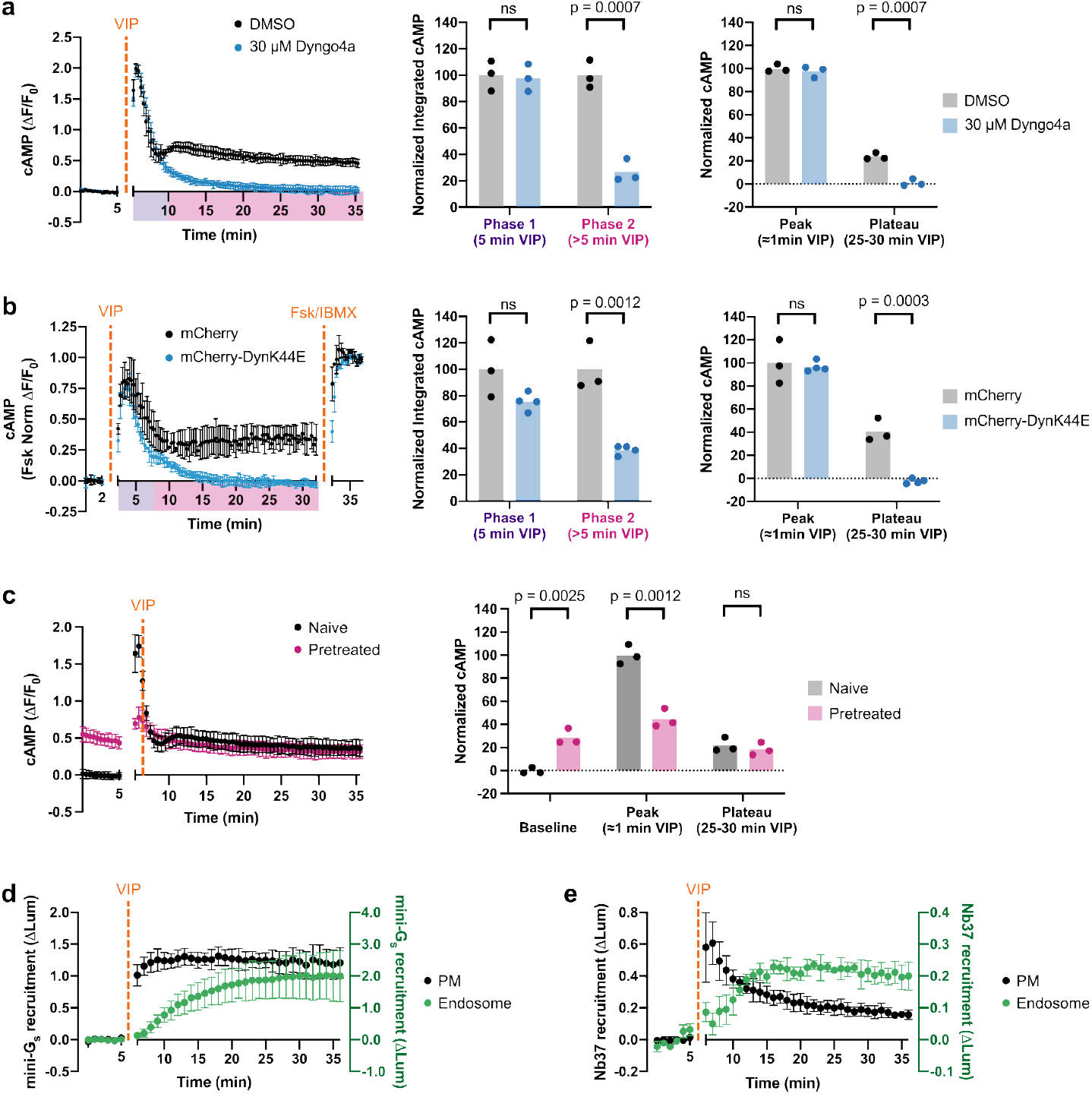
cAMP signaling through endogenous VIPR1 is biphasic. **a**, Intracellular cAMP, as measured by the fluorescence change of cADDis cAMP biosensor, upon treatment with 500 nM VIP added at 5 minutes. Cells were pretreated with 30 µM Dyngo4a (blue, n = 4) or DMSO (black, n = 3) for 10 minutes. Integrated cAMP of two phases (0-5 min and 5-30 min VIP treatment) was calculated as the area under the curve and normalized to the average DMSO value. Normalized changes in fluorescence for the peak (maximum ΔF/F_0_) and the plateau (average for timepoints 25-30 min VIP treatment) were normalized to the average DMSO peak value. Significance was determined by unpaired t-tests. **b**, Changes in cAMP in cells expressing mCherry-Dyn1K44E (blue, n = 4) or mCherry (black, n = 3) upon treatment with 1 μM VIP added at 2 minutes. Fluorescence was measured by microscopy and normalized to fluorescence change upon co-application of forskolin (Fsk, 10 µM) and IBMX (500 µM) at 32 minutes. Integrated cAMP and peak/plateau ΔF/F_0_ were quantified as in (**a**), with data normalized to the mCherry average. **c**, Changes in cAMP in cells pretreated with VIP (“Pretreated,” pink) compared to those not (“Naive,” black). Pretreated samples were treated with 500 nM VIP for 10 minutes, followed by a 10 minute washout period prior to the acquisition of baseline measurements. Baseline (average for 5 min baseline), peak, and plateau ΔF/F_0_ were quantified as in (**a**), with data normalized to the average naive peak value. n = 3. **d-e**, NanoBiT bystander assays showing recruitment of mini-G_s_ (**d**, n = 5) or Nb37 (**e**, n = 3) to the plasma membrane (black) or the endosome (green) upon addition of 1 µM VIP at 5 minutes. For all panels, data represent biological replicates and are shown as individual data points or mean ± s.d.

As VIPR1 internalization does not require β-arrestin, we wanted to confirm that it indeed recruits β-arrestin after activation by VIP^31^. Live confocal fluorescence imaging demonstrated visible membrane recruitment of β-arr1 after VIP application, first to the plasma membrane and then to endosomes containing internalized VIPR1 (Fig. 1e). We further quantified this sequential β-arrestin recruitment phenotype with a nanoluciferase protein complementation (NanoBiT) assay using CAAX motif or endofin-derived FYVE domain constructs as previously validated markers of the plasma membrane and endosome membrane, respectively (Extended Data Fig. 3)^35^. Using this ‘bystander’ assay, we observed sequential recruitment of β-arr2 after VIP-induced activation of VIPR1, first to the plasma membrane and then to the endosome membrane, with the transition between the two being nearly complete within five minutes (Fig. 1f).

**Fig. 3.**
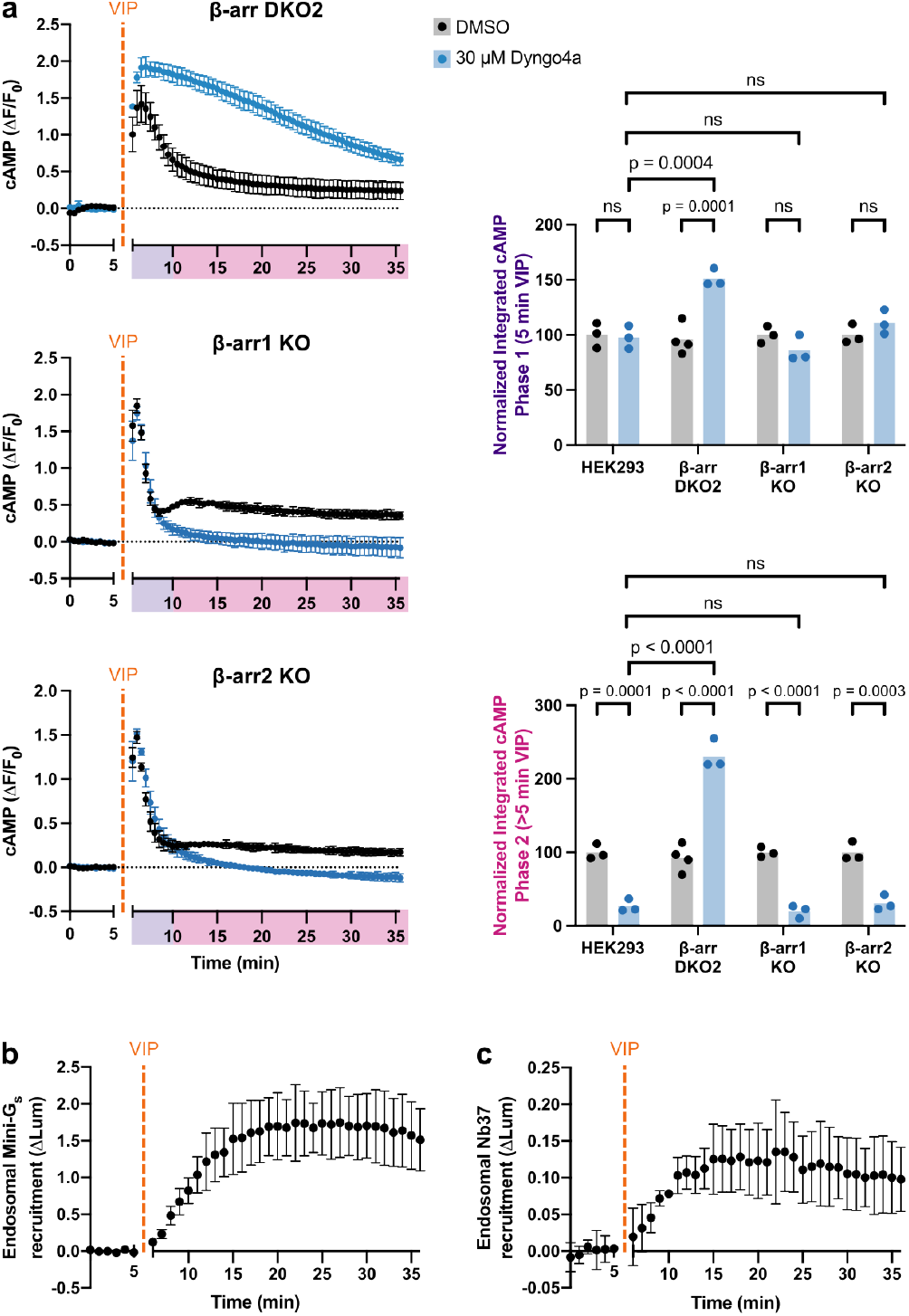
β-arrestin is dispensable for VIPR1 endosomal cAMP signaling. **a**, Changes in cAMP in β-arr1/2 DKO line (β-arr DKO2) and β-arr1/2 single KO lines (β-arr1 KO and β-arr2 KO) upon treatment with 500 nM VIP added at 5 minutes. Cells were pretreated with 30 µM Dyngo4a (blue) or DMSO (black) for 10 minutes. Integrated cAMP of each phase (0-5 min or 5-30 min VIP treatment) was calculated as the area under the curve and normalized to the average DMSO value for each cell line. Data for WT HEK293 repeated from **Fig. 2a**. Significance was determined by a two-way ANOVA with Tukey’s multiple comparisons test along with data for β-arr DKO1 cell line shown in **Extended Data Fig. 5a**. n = 3-4, as indicated. **b-c**, NanoBiT bystander assays in β-arr DKO2 cells showing endosomal recruitment of mini-G_s_ (**b**, n = 3) or Nb37 (**c**, n = 4) upon addition of 1 µM VIP at 5 minutes. For all panels, data represent biological replicates and are shown as individual data points or mean ± s.d.

### VIPR1 mediates sequential phases of cAMP production from the plasma membrane and endosomes

These above results indicate that VIPR1 robustly recruits β-arrestin and remains bound to β-arrestin in endosomes, yet it does not require β-arrestin for endocytosis. To our knowledge, these characteristics differentiate VIPR1 from all other GPCRs for which the β-arrestin-dependence of endosomal signaling has been explicitly investigated. This motivated us to ask if VIPR1 is able to generate an endocytosis-dependent cAMP signaling phase.

VIPR1 is natively expressed in the kidney^36^, and previous studies have documented endogenous VIPR1 expression in HEK293 cells^31,37,38^. We first sought to independently verify that endogenous VIPR1 is indeed responsible for the VIP-elicited cAMP elevation measured in our cell model. CRISPR-mediated knockout of VIPR1 strongly suppressed the cytoplasmic cAMP elevation elicited by VIP application, as measured in living cells using a genetically encoded fluorescent cAMP biosensor (cADDis), while VIPR2 knockout had little effect (Extended Data Fig. 4a-b). A small residual cAMP response was still present in VIPR1 knockout cells, and other secretin family GPCRs--notably the secretin and pituitary adenylate cyclase-activating polypeptide (PACAP) receptors--are also sensitive to VIP^39,40^. The PACAP 1 receptor (PAC1R) agonist PACAP-27 produced a detectable cAMP response exceeding that produced by VIP in VIPR1 KO cells, while secretin had no detectable effect (Extended Data Fig. 4c). These observations suggest that the residual VIP response is mediated by low-level expression of PAC1R which displays a ∼100-fold lower EC_50_ for PACAP-27 versus VIP (Extended Data Fig. 4d)^40^ and has been detected in HEK293 by RNA-Seq^38^. However, as this residual contribution to the cellular cAMP elevation is very small when compared to the VIPR1-mediated response (Extended Data Fig. 4b), we conclude that VIPR1 is indeed the major endogenous GPCR mediating the VIP-induced cAMP elevation measured in this study.

**Fig. 4.**
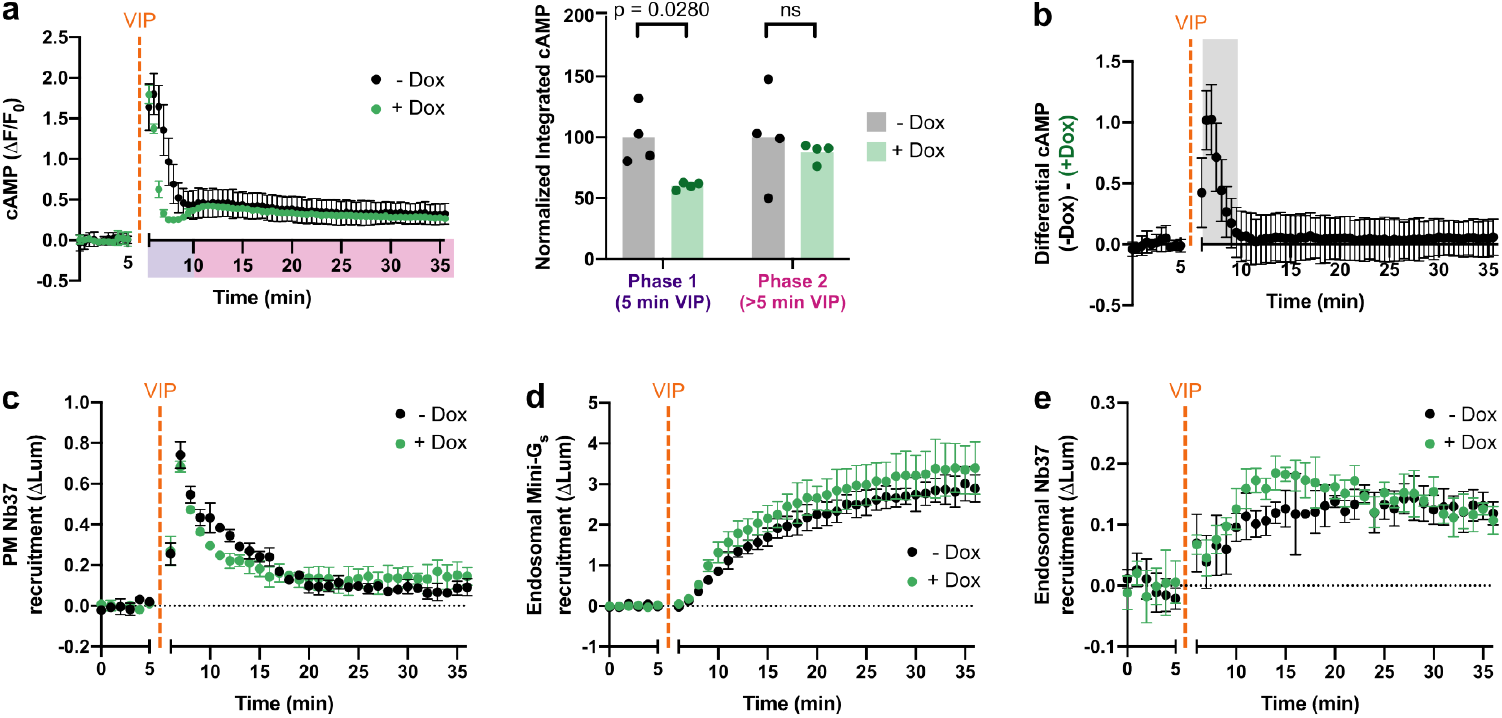
β-arrestin resolves phases of cAMP signaling. **a**, Changes in cAMP in β-arr DKO2-R with (green) and without (black) Dox induction upon treatment with 500 nM VIP added at 5 minutes. Integrated cAMP of each phase (0-5 min or 5-30 min VIP treatment) was calculated as the area under the curve and normalized to the average -Dox value. Significance was determined by a repeated measures 2 way ANOVA with Sidak’s multiple comparisons test along with data for DKO1-R shown in Extended Data Fig. 7a. n = 4. **b**, Difference in cAMP response with and without β-arr2 expression, calculated as the difference between the -Dox and +Dox curves shown in (**a**). Shaded area represents timepoints where the difference is statistically significant, as determined by a repeated measures 2 way ANOVA (p < 0.05) with Sidak’s multiple comparisons test carried out on the curves in (**a**). **c-e**, NanoBiT bystander assays showing plasma membrane recruitment of Nb37 (**c**, n = 3) or endosomal recruitment of mini-G_s_ (**d**, n = 3) or Nb37 (**e**, n = 4) in β-arr DKO2-R cells upon addition of 1 µM VIP at 5 minutes. For all panels, cells treated with Dox to induce expression of β-arr2-mApple are shown in green, and data represent biological replicates, shown as individual data points or mean ± s.d.

Next, we characterized the time course of the VIP-induced cAMP signal, as defined for these experiments by elevation of global cAMP concentration in living cells at 37°C. Stimulation of HEK293 cells with VIP elicited a complex signal, characterized by an initial phase of cAMP elevation that peaked within <1 minute and desensitized over ∼5 minutes followed by a second, less intense peak and a later plateau of cAMP elevation that persisted above baseline in the prolonged presence of VIP (Fig. 2a, black curve). Both of these signaling phases were lost in VIPR1 knockout cells, indicating that both are mediated by VIPR1 (Extended Data Fig. 4a). Furthermore, analysis of cAMP dynamics using quantitative fluorescence microscopy detected both phases in individual cells (Extended Data Fig. 4e). Accordingly, both phases of the complex VIP-induced cellular cAMP signal occur in the same cells and are mediated by endogenous VIPR1.

To investigate how endocytic trafficking impacts this overall cellular cAMP signal, we examined the effect of experimentally imposing endocytic blockade on the global cytoplasmic cAMP elevation. Preventing dynamin-dependent endocytosis of VIPR1 using a chemical inhibitor, Dyngo4a, abolished the second cAMP peak and subsequent plateau elevation without detectably affecting the first peak (Fig. 2a., blue curve, and Extended Data Table 1). Genetically imposing endocytic inhibition with Dyn1-K44E also selectively blocked the second phase (Fig. 2b). These results are consistent with the VIPR1-mediated cellular cAMP elevation being composed of two sequential phases: a first phase from the plasma membrane producing an initial peak of glocal cAMP elevation, and then a second phase from endosomes producing a subsequent peak and plateau. We further noted that cells pretreated with VIP exhibited an elevated baseline cAMP that persisted after VIP washout, whereas rechallenge with VIP was unable to produce a subsequent additional elevation (Fig. 2c). These results suggest that endosomal cAMP signaling by VIPR1 can be sustained even after agonist removal, despite the plasma membrane signal being terminated and desensitized within several minutes.

To further validate the subcellular location(s) of ligand-dependent VIPR1 and G_s_ activation, we took advantage of conformational biosensors that detect either active-state VIPR1 (mini-G_s_)^41,42^ or a conformational intermediate in the process of G_s_ activation (Nb37)^12,43^. Using these tools in NanoBiT bystander assays, we observed that VIP promoted the recruitment of both biosensors first to the plasma membrane and then to endosomes (Fig. 2d,e). These results suggest that VIP-induced activation of VIPR1 and G_s_ occurs at both membrane locations, sequentially and with kinetics consistent with the biphasic cytoplasmic cAMP elevation measured using the cAMP biosensor.

### β-arrestin and endocytosis both promote VIPR1 desensitization at the plasma membrane, but β-arrestin is dispensable for the second endosomal phase of VIPR1-mediated cAMP signaling

Having established the unique spatial and temporal profile of VIPR1-mediated cAMP signaling, we next asked how β-arrestin regulates it given that VIPR1 trafficking is β-arrestin-independent. To investigate this, we first compared endogenous VIPR1-mediated cAMP signaling produced in β-arr DKO cells to that produced in the parental (wild type) cell background. In β-arr DKO cells, VIP elicited a rapid cAMP peak followed by a decay (Fig. 3a and Extended Data Fig. 5a). Calculating first derivatives and fitting decay time constants to these curves indicated that the initial cAMP peak observed in β-arr DKO cells is slightly more prolonged when compared to that observed in parental wild type cells (Extended Data Fig 5b, Extended Data Table 1). Nevertheless, the initial cAMP peak measured in β-arr DKO cells still obviously decayed, with β-arr1/2 depletion slowing the decay time constant only by ∼2-fold. This indicates that a second, β-arrestin-independent mechanism for terminating the initial VIPR1 signaling peak must exist. We found that imposing endocytic blockade with Dyngo4a dramatically prolonged the VIP-induced cAMP elevation measured in β-arr DKO cells (Fig. 3a, Extended Data Fig. 5a, Extended Data Table 1), in contrast to its much smaller effect in wild type cells. This indicates that endocytosis by itself can terminate the initial VIPR1-mediated cAMP elevation in cells lacking β-arrestin. That endocytosis itself can attenuate the first cAMP peak without β-arrestin is fully consistent with β-arrestin-independent internalization of VIPR1, and it suggests that endocytosis and β-arrestin semi-redundantly terminate the first cAMP peak in wild type cells.

**Fig. 5.**
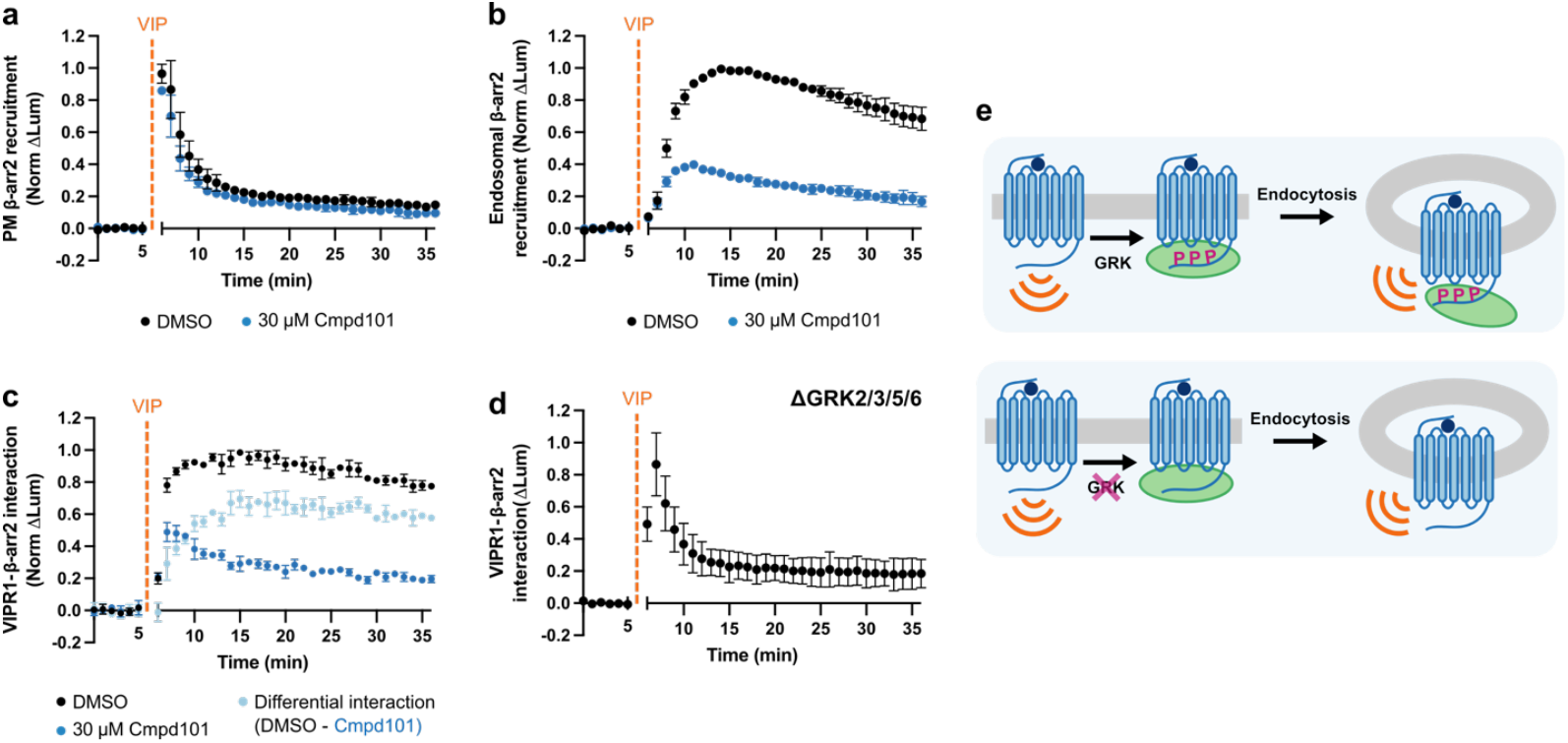
VIPR1-β-arrestin complexes at the plasma membrane and endosomes are differentially regulated by GRKs. **a-b**, NanoBiT bystander assays showing plasma membrane (**a**) or endosomal (**b**) recruitment of β-arr2 in HEK293 cells upon addition of 1 µM VIP at 5 minutes. Cells were pretreated with DMSO (black) or 30 µM Cmpd101 (blue) for 10 minutes. **c-d**, Direct NanoBiT assay showing recruitment of β-arr2 to VIPR1 in HEK293 (**c**) or HEK293A GRK2/3/5/6 knockout cells (**d**) upon addition of 1 µM VIP at 5 minutes. HEK293 cells were pretreated with DMSO (black) or 30 µM Cmpd101 (blue) for 10 minutes. Light blue curve in (**c**) represents the difference in interaction over time due to Cmpd101, calculated as the difference between the DMSO and Cmpd101 curves. **e**, Model for GRK regulation of VIPR1-β-arrestin complexes. Upon VIP stimulation, VIPR1 is phosphorylated by GRKs, leading to the recruitment of β-arrestin to VIPR1 at both the plasma membrane and endosomes. If VIPR1 phosphorylation by GRKs is blocked, β-arrestin is still recruited to VIPR1 at the plasma membrane, but a VIPR1-β-arrestin complex does not form at the endosome. For all panels, data represent three biological replicates shown as mean ± s.d.

β-arr1 and β-arr2 are both endogenously expressed in HEK293 cells^38^ and, while we observed similar recruitment of each isoform to activated VIPR1 (Fig. 1e,f), we wondered whether the signal-attenuating effect might be unique to one isoform. To explore this question, we measured VIP-induced cAMP signaling in β-arrestin single knockout cell lines (Extended Data Fig. 5c, d). In cells selectively deficient in β-arr1 or β-arr2, VIP application elicited cAMP elevation similar in kinetics to that observed in wild type cells (Fig. 3a, Extended Data Table 1). Importantly, in both single KO cell backgrounds, VIPR1 desensitization was observed upon endocytic inhibition, and blocking endocytosis reduced the second phase of signaling to a degree comparable to the reduction observed in wild type cells (Fig. 3a). Therefore, both β-arrestins appear similar in their overall effects on VIPR1 cAMP signaling as assessed with the present methods.

While we did not observe a distinct, second cAMP peak in β-arr DKO cells, a plateau above baseline was still observed (Fig. 3a). This contrasts with a complete loss of the cAMP plateau produced by endocytic inhibition in wild type cells (Fig. 2a, 3a, Extended Data Fig. 5a). Further, VIP-induced endosomal recruitment of both mini-G_s_ and Nb37 was still observed in β-arr DKO cells (Fig. 3b,c, Extended Data Fig. 5e,f). Together, these results indicate that β-arrestin is not required for VIPR1 to initiate either the plasma membrane or endosomal phases of G_s_ activation or cAMP elevation.

### A discrete role of β-arrestin in temporally resolving the VIPR1-mediated signaling phases

If β-arrestin binds active VIPR1 but is not necessary for receptor trafficking or signaling, then what role does it play in modulating VIPR1 activity? That the β-arr DKO cell lines displayed a broader, Dyngo4a-sensitive cAMP peak compared to the sharply biphasic cAMP production in parental cells suggests β-arrestin sculpts response kinetics. To more precisely investigate this, we pursued genetic rescue in β-arr DKO cells using recombinant β-arr2 (β-arr2-mApple) expressed under tetracycline-inducible control (DKO-R, Extended Data Fig. 6). As expected, a comparison of real-time cAMP dynamics in these cells indicated that overexpression of β-arr2 (+Dox) accelerates the desensitization of the first phase of VIP-induced cytoplasmic cAMP elevation relative to the uninduced (-Dox) condition (Fig. 4a,b and Extended Data Fig. 7a,b). We further quantified this effect by fitting decay constants to the desensitization of the first phase, which showed around a two-fold increase upon β-arr2 induction (Extended Data Table 1), as well as by calculating the first derivative of the curves, which became more negative upon β-arr2 induction (Extended Data Fig. 7c). While we observed a subtle corresponding decrease in Nb37 recruitment to the plasma membrane upon β-arr2 induction, this trend was not statistically significant when assessed across the two independent β-arr DKO cell lines (Fig. 4c and Extended Data Fig. 7d).

**Fig. 6.**
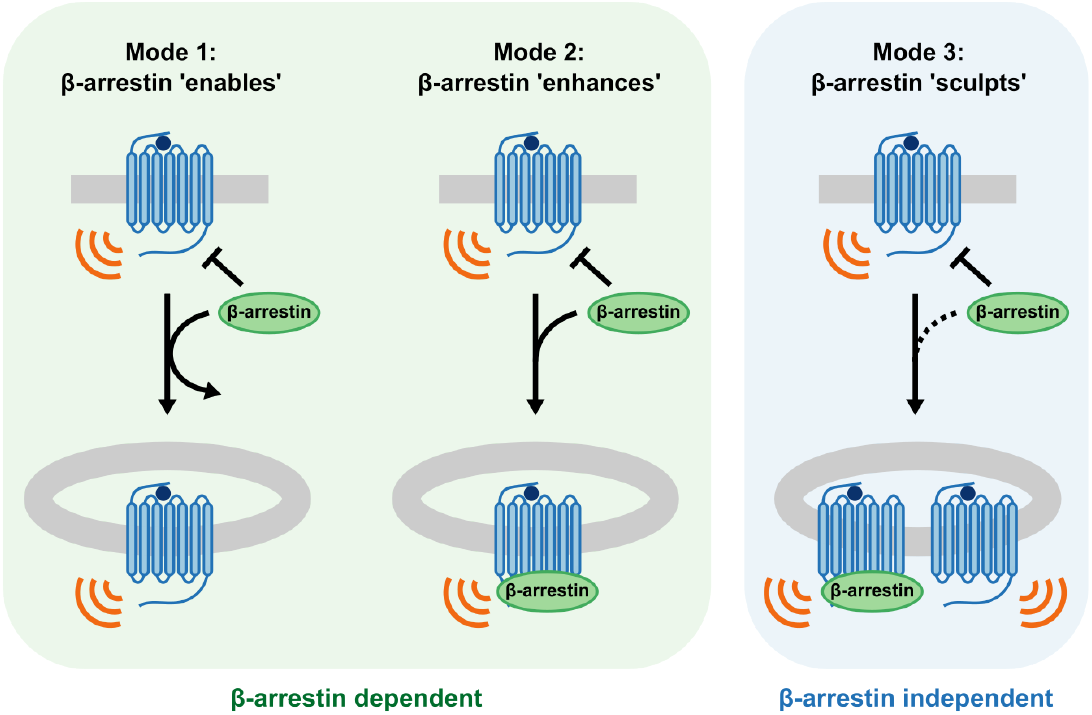
VIPR1 defines a third mode of endosomal GPCR signaling. Diagram illustrates key distinctions among signaling modes. *Mode 1:* β-arrestin enables signaling from endosomes by promoting receptor internalization and then dissociating. *Mode 2:* β-arrestin promotes receptor internalization and then remains bound after internalization to form a signal-sustaining endosomal megaplex. *Mode 3:* β-arrestin is not required for receptor internalization or signaling from endosomes; instead, β-arrestin sculpts the resulting combined signal output, such that the plasma membrane and endosomal signal phases produce temporally resolved cAMP peaks.

Some GPCRs that bind β-arrestin in endosomes use this interaction to potentiate cAMP signaling^2,11,24–28^. However, the cellular cAMP elevation observed during the plateau phase of VIPR1 signaling was not detectably affected by β-arr2 overexpression, and the only statistically significant difference in the cAMP elevation with or without β-arr2 overexpression occurred in the first 5 minutes after VIP application (Fig. 4a,b and Extended Data Fig. 7a,b). This observation indicates that β-arrestin, despite attenuating the plasma membrane signaling phase, does not detectably modulate the endosomal cAMP signaling phase either positively or negatively. We returned to the NanoBiT bystander assays to explore this further. In agreement with the bulk cAMP assays, neither mini-G_s_ nor Nb37 recruitment to the endosome was markedly altered by β-arr2 overexpression in the two β-arr DKO cell lines tested (Fig. 4d,e, Extended Data Fig. 7e,f).

Together, these data verify that β-arrestin is not necessary for VIPR1-mediated signaling from endosomes and indicate that β-arrestin binding differentially modulates VIPR1 signaling activity depending on the subcellular location of the complex. The net effect of this selective modulation, with β-arrestin complex formation at the plasma membrane being signal-attenuating and at endosomes being signaling-neutral, is to cause the plasma membrane and endosomal signaling phases to resolve as separate cAMP peaks. Accordingly, β-arrestin’s primary function in VIPR1 signaling appears to be to convert distinct phases of VIPR1-Gs activation that are initiated from spatially resolved subcellular locations into temporally resolved phases of global cytoplasmic cAMP elevation.

### GRK-mediated phosphorylation of VIPR1 differentially regulates β-arrestin binding at the plasma membrane and endosomes

Our data using the β-arr DKO rescue cell lines indicate that VIPR1-β-arrestin complexes are functionally different depending on their subcellular localization. To begin to investigate the biochemical underpinnings of these functional differences, we explored how GRK-mediated phosphorylation of VIPR1 affects complex formation at the plasma membrane and endosomes. Chemical inhibition of GRK2/3 in HEK293 cells using Compound 101 (Cmpd101)^44^ did not significantly affect VIP-stimulated recruitment of β-arr2 to the plasma membrane, but it strongly inhibited recruitment to endosomes (Fig. 5a, b). This differential effect of Cmpd101 on β-arrestin recruitment elicited by VIPR1 at each membrane location stands in contrast to recruitment of β-arrestin by V2R, a GPCR that also binds β-arrestin at both locations^11,45^. For the V2R, Cmpd101 markedly inhibited β-arr2 recruitment at both the plasma membrane and endosomes (Extended Data Fig. 8a,b). The selective effect on endosomal recruitment of β-arrestin was not due to a lack of VIPR1 internalization, as Cmpd101 had no effect on the agonist-induced endocytosis of VIPR1 (Extended Data Fig. 8c). Further, a direct NanoBiT assay examining the time course of VIPR1 / β-arr2 binding indicated that complex sensitivity to Cmpd101 is itself location-dependent, with observed inhibition increasing at later time points (>5 min VIP) that correspond to when receptors are present in endosomes (Fig. 5c). Providing genetic confirmation of this pharmacological result, VIPR1 / β-arr2 complexes were rapidly formed but transient in ΔGRK2/3/5/6 cells^46^ (Fig. 5d). Together, these results strongly suggest that VIPR1 / β-arrestin complexes formed at the plasma membrane and endosomes are biochemically as well as functionally distinct. The VIPR1 / β-arrestin complex which terminates signaling at the plasma membrane does not require GRK-mediated phosphorylation of receptors to form. The complex present on the endosome limiting membrane that is signaling-neutral, in marked contact, is specifically GRK-dependent (Fig. 5e).

## Discussion

The present findings take advantage of the distinct trafficking properties of VIPR1 to disentangle the effects of β-arrestin and endocytosis on the G_s_-coupled cellular cAMP response elicited by this GPCR. By doing so, our results establish VIPR1 as a GPCR that is capable of internalizing and initiating a second phase of endosomal G_s_-cAMP signaling in the complete absence of β-arrestin. We further show that β-arrestin plays a different role in determining the cellular VIPR1 response by resolving the plasma membrane and endosome signaling phases into separate and sequential global cAMP peaks. We note that a number of GPCRs internalize in a β-arrestin-independent manner^23,30^, and some, such as the glucagon-like peptide-1 receptor (GLP1R), have also separately been shown to signal from endosomes^14^. However, to our knowledge, the present results are the first to explicitly show, for any GPCR, generation of an endosomal G_s_-cAMP signal in the complete absence of cellular β-arrestin.

Accordingly, we propose to revise the present concept of endosomal signaling by GPCRs and, in particular, expand the understanding of how receptor interactions with β-arrestin impact the spatiotemporal profile of cAMP signaling (Fig. 6). For receptors that weakly bind β-arrestin, exemplified by the thyroid stimulating hormone receptor (TSHR), β-arrestin *enables* signaling to occur from endosomes by promoting receptor internalization before dissociating^9,47^. For some receptors that more strongly bind β-arrestin, exemplified by PTHR1 and V2R, β-arrestin *enhances* the endosomal signal by remaining bound to form a signal-boosting complex specifically on the endosome membrane ^2,24–26,28^. In the case of VIPR1, β-arrestin is not required to generate the endosomal signal. Rather, it *sculpts* receptor activity such that sequential peaks of receptor / G protein activation initiated from different subcellular locations – the plasma membrane and endosomes – are resolved into temporally separated peaks of global cytoplasmic cAMP elevation. As β-arrestin is not needed even to deliver receptors to the endosome, this third mode of endosomal signaling reveals a discrete function of β-arrestin in regulating the spatiotemporal profile of the integrated cellular cAMP signal. Because other GPCRs have been reported to internalize independently of β-arrestin^23,30^, we anticipate that the additional endosomal signaling mode and β-arrestin function revealed in the present study is not unique to VIPR1 and likely underlies GPCR-specific modulation of cellular signaling more broadly.

β-arrestin mediates this signal-sculpting function by selectively accelerating the rate at which the plasma membrane signaling phase is attenuated while having little or no effect on the endosomal signaling phase. The mechanistic basis for this specificity in β-arrestin function appears to be the presence of distinct GPCR / β-arrestin complexes at the plasma membrane relative to at the endosome limiting membrane, as established here by differences in their dependence on GRK-mediated receptor phosphorylation for formation or stability. This supports the emerging view that GPCR / β-arrestin complexes dynamically remodel during endocytic trafficking to enable location-specific control^25,28,29,48,49^. Further investigation will be needed to determine the biophysical basis for this biochemical distinction. Transition from a ‘core engaged’ GPCR / β-arrestin complex to a ‘tail engaged’ complex is thought to underlie the ability of β-arrestin to switch from a signal-attenuating factor at the plasma membrane to a signal-enhancing factor at the endosome for receptors such as PTHR^25,28,29^. Therefore, a simple model is that the signal-attenuating VIPR1 / β-arrestin complex formed at the plasma membrane corresponds to a receptor ‘core-engaged’ complex that can form in the absence of a phosphorylated receptor tail, while the signaling-neutral VIPR1 / β-arrestin complex formed at endosomes corresponds to a phosphorylation-dependent ‘tail-engaged’ complex (Fig 5e). While there are examples of signaling neutral GPCR-β-arrestin mutant complexes that are permissive of G protein signaling without modulating the cAMP response^28,50^, it remains unclear how a signaling-neutral VIPR1-β-arrestin complex may be distinct from previously described ‘tail engaged’ signal-enhancing GPCR-β-arrestin complexes. Thus, we anticipate that additional diversity exists in GPCR / β-arrestin complex formation to enable the functional signaling properties of GPCRs to be precisely tuned in a receptor-specific manner, and according to differences in the post-translational modification of receptors at discrete membrane locations.

## Methods

### Cell culture

HEK293 (ATCC CRL-1573) and HEK239A ΔGRK2/3/5/6^46^ were cultured in Dulbecco’s modified Eagle’s medium (DMEM, Gibco 1196511) supplemented with 10% FBS (Hyclone SH30910.03). Besides ΔGRK2/3/5/6, all other knockout and stable cell lines used in this study were derived from the HEK293 cell line. Cells stably expressing FLAG-VIPR1 (CMV) were selected using 500 μg/mL geneticin, and cells stably expressing HaloTag-VIPR1 (CMV) or HaloTag-β2AR were selected using 25 μg/mL zeocin. Cell lines stably expressing tet-inducible (TRE3G) HaloTag-VIPR1 or β-arr2-mApple were selected using 2 µg/mL puromycin and screened for inducible expression by flow cytometry. Rescue was induced 24 hours before experiments using 1 ug/mL doxycycline. All cells were routinely checked for mycoplasma contamination (MycoAlert, Lonza).

### DNA constructs

Information on all plasmids used in this study can be found in Supplementary Table 1. Lipofectamine 2000 (Thermo Fisher) was used for transfections according to the manufacturer’s protocol. BacMam from pCMV-Dest constructs were produced according to the manufacturer’s protocol.

### Generation of CRISPR knockout cell lines

sgRNAs (Supplementary Table 2) were designed using the Synthego CRISPR design tool. To make RNP, 3 µL total of 53.3 µM sgRNA (Synthego) was mixed with 2 µL of 40 µM Cas9 (UC Berkeley Macrolab), and the mixture was incubated at room temperature for 10 minutes. Electroporation of the RNP complex was carried out using the SF Cell Line 4D-Nucleofector kit (Lonza) and program CM-130 with 2.0×10^5^ cells. Genetic modifications of monoclonal cell lines were verified from genomic PCR amplicons (Supplementary Table 2-3) using Sanger sequencing in combination with the Synthego ICE Analysis Tool or by next generation sequencing (Amplicon-EZ, Azenta Life Sciences). β-arr1/2 expression was assessed by western blot (Supplementary Table 4) quantified using Fiji^51^. For the β-arr DKO1 cell line, ARRB1 and ARRB2 modifications were done simultaneously, while for the β-arr DKO2 cell line, modifications were done sequentially.

### cADDis microplate cAMP assay

Real-time intracellular cAMP was measured using the Green Up cADDis cAMP biosensor (Montana Molecular), as described previously^52^. Briefly, cells were treated with cADDis BacMam and plated into 96-well plates (Corning 3340). After 24 hours, cells were washed with assay buffer (20 mM HEPES pH 7.4, 135 mM NaCl, 5 mM KCl, 0.4 mM MgCl_2_, 1.8 mM CaCl_2_, 5 mM d-glucose) twice before a 10 minute incubation in a 37°C plate reader (Synergy H4, BioTek), optionally with DMSO or Dyngo4a (30 μM, Abcam ab120689) as indicated in figure legends. Fluorescence was read at an excitation wavelength of 500 nm and an emission wavelength of 530 nm every 20-30 seconds for 35 minutes, with vehicle or agonist added after 5 minutes. Change in intracellular cAMP (ΔF / F_0_) was calculated as the change in fluorescence (ΔF = F - F_0_) normalized to the average of the 5 minute baseline (F_0_). For dose-response curves, peak cAMP values, normalized to the average peak response to 1 µM PACAP, were fit to the equation: 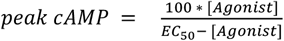

### cADDis live epifluorescence microscopy

Cells were transfected with the appropriate constructs 48 hours prior to experiments and plated into glass bottom dishes (MatTek Corporation) coated with 0.001% (w/v) poly-L-lysine (Millipore Sigma). Cells were washed three times with assay media (DMEM, no phenol red, 30 mM HEPES pH 7.4) before imaging. Imaging was conducted using a Nikon TE2000 inverted microscope fitted with a Nikon 20x/0.7 NA objective, Sutter Lambda HPX-L5 LED epifluorescence illuminator, Sutter excitation / emission filter wheels and Point Grey CMOS camera controlled by Micromanager^53^ (www.micro-manager.org). A dual channel dichroic filter set (Chroma 89021) was used to separately acquire cADDis (470/40 excitation, 520/50 emission) and mCherry (572/35 excitation, 632/60 emission) fluorescence signals. Cells were maintained at 37°C during image collection using a custom-built heated stage and stage top incubator. 1 µM VIP was added after a 2 minute baseline, and forskolin (Fsk, 10 µM, MilliporeSigma) and 3-isobutyl-1-methylxanthine (IBMX, 500 µM, MilliporeSigma) were added 30 minutes later. Data was processed using Cell Profiler 4^54^ (www.cellprofiler.org). Briefly, cells were identified using the mCherry channel, and the intensity of the cADDis channel was measured over the entire image and in the cell mask. Each frame was background corrected by subtracting the lower quartile fluorescence intensity of the entire image. The normalized change in fluorescence of the cell mask was calculated as (F – F_0_)/(F_FskIBMX_ - F_0_), where F_0_ is the average fluorescence over the first two minutes and F_FskIBMX_ is the average fluorescence of the final two minutes. Each biological replicate is the average of 1-2 dishes of approximately 20-30 cells per dish.

### NanoBiT luciferase complementation assays

Cells grown in 6-well plates were transfected with receptor (HaloTag-VIPR1 or FLAG-V2R) and the appropriate LgBit/SmBiT constructs 24 hours prior to experiments. Experiments using Nb37-SmBiT(114) (Fig. 4c, Extended Data Fig. 7d) or Nb37-SmBiT(101) (all other figures) also required the cotransfection of a tricistronic G protein construct (human G?1, bovine Gγ1, and human Gαs). For the assay, cells were lifted and centrifuged at 500 x g for 3 minutes. Cells were resuspended in assay buffer supplemented with 5 µM coelenterazine-H (Research Products International) to a concentration of 5×10^5^ or 1×10^6^ cells/mL, and 100 µL of cells were plated into untreated white 96-well plates (Corning 3912). Plates were incubated at 37 °C and 5% CO_2_ for 10 minutes prior to the assay, optionally with DMSO or Cmpd101 (30 μM, HelloBio HB2840) as indicated in figure legends. Luminescence was read on a plate reader (Synergy H4, BioTek) prewarmed to 37 °C. After a 5 minute baseline reading, vehicle or agonist were added at concentrations indicated in figure legends, and luminescence was read for another 30 minutes. To calculate the change in luminescence, each well was first normalized to its average baseline luminescence, and the average change in luminescence for vehicle-treated samples was subtracted from the average change in luminescence for drug-treated samples (ΔLum = Lum_VIP_ - Lum_vehicle_). To compare DMSO- and Cmpd101-treated samples, readings were further normalized to the maximum ΔLum of the DMSO-treated sample (Norm ΔLum).

### Confocal microscopy

All fixed and live confocal imaging was carried out using a Nikon Ti inverted microscope fitted with a CSU-22 spinning disk unit (Yokogawa), custom laser launch (100 mW at 405, 488, 561, and 640 nm, Coherent OBIS), Sutter emission filter wheel and Photometrics Evolve Delta EMCCD camera. Samples were imaged using an Apo TIRF 100x/1.49 NA oil objective (Nikon).

For live imaging, cells were transfected with the appropriate constructs 48 hours prior to experiments and plated into glass bottom dishes (Cellvis) coated with 0.001% (w/v) poly-L-lysine (Millipore Sigma). For surface labeling of receptor, cells were incubated with either monoclonal anti-FLAG M1 antibody (Millipore Sigma F3040) labeled with Alexa Fluor 647 (Thermo Fisher A20186) or 200 nM JF_635_i-HTL^32^ for 10 minutes at 37 °C and 5% CO_2_. After three washes, cells were imaged in assay media (DMEM, no phenol red, 30 mM HEPES pH 7.4) in a temperature- and humidity-controlled chamber (Okolab). For time-lapse, cells were imaged at 20 second intervals for 22 minutes, with 500 nM VIP added after 2 minutes.

For fixed imaging, cells were transfected with the appropriate constructs 48 hours prior to experiments and plated onto coverslips (Fisher Scientific 12-545-100P) coated with 0.001% (w/v) poly-L-lysine (Millipore Sigma). Coverslips were washed with assay media (DMEM, no phenol red, 30 mM HEPES pH 7.4) and labeled with 200 nM JF_635_i-HTL for 10 minutes at 37 °C and 5% CO_2_. Vehicle or 500 nM VIP was added for 20 minutes, followed by washes with PBS and fixation in 3.7% formaldehyde (Fisher, F79) in modified BRB80 (80 mM PIPES pH 6.8, 1 mM MgCl2, 1 mM CaCl2) for 20 minutes. For immunofluorescence, cells were permeabilized with 0.1% Triton-×100 in 3% BSA before incubation with primary and secondary antibodies (Supplementary Table 4). Coverslips were mounted on glass slides using ProLong Gold Antifade mounting media (Invitrogen P10144).

All image analysis was carried out in Cell Profiler 4^54^, and all images for figures were processed using Fiji^51^. For colocalization analysis, the Pearson’s correlation coefficient was calculated in individual cells segmented manually.

### Receptor internalization by flow cytometry

All experiments were carried out using cell lines stably expressing an N-terminal HaloTag fusion of the receptor of interest. Unless otherwise noted, all experiments use HaloTag-VIPR1 under the control of an inducible promoter (TRE3G) in either a VIPR1 KO or β-arr1/2 DKO cell background. For experiments using mCherry and mCherry-Dyn1K44E, cells were incubated with BacMam for 24 hours prior to the experiment. Agonist or vehicle was added to cells grown in 12-well plates coated with 0.001% (w/v) poly-L-lysine (Millipore Sigma) and incubated at 37 °C and 5% CO_2_ as indicated in figure legends. At the end of the incubation, plates were cooled on ice for 10 minutes, and all subsequent steps were carried out on ice. Cells were then labeled with 200 nM JF_635_i-HTL for 30 minutes. After washing once with PBS-EDTA (UCSF Cell Culture Facility), cells were lifted with 100 uL of TrypLE Express (Thermo Fisher) at room temperature for 10 minutes. TrypLE was quenched with 150 uL PBS-EDTA + 2.5% FBS, and resuspended cells were transferred into an untreated black 96-well plate (Corning). Data from 10,000 events was collected on an Attune NxT Flow Cytometer equipped with a CytKick Autosampler (Thermo Fisher). JF_635_i-HTL was measured with a 637 nm excitation laser and a 670/14 nm emission filter, while mCherry was measured with a 561 nm excitation laser and a 620/15 nm emission filter. Data were analyzed using FlowJo v10.8 Software (BD Life Sciences). Populations were gated for cells expressing mCherry, when applicable (Supplementary Figure 1). Surface receptor was calculated as the normalized median fluorescence (JF_635_i).

### Statistical analysis and reproducibility

All data are shown as individual biological replicates or as a mean ± standard deviation from at least 3 biologically independent experiments. Each biological replicate in the cAMP, NanoBiT, and flow cytometry assays represents the average of at least two technical replicates, and all images are representative of at least 3 biologically independent experiments. Rate constants, derivatives, and statistical tests were carried out using Prism 8/9 (GraphPad) as noted in figure legends.

## Acknowledgements

We thank D. Gordon and A. Ehrlich for assistance with CRISPR; L. Lavis for helpful discussion and sharing reagents; A. Inoue, G. Schulte, R. Irannejad, and A. Marley for sharing reagents; B. Barsi-Rhyne and the rest of the von Zastrow lab for helpful discussion; and the following core facilities for providing services to support this research: the UCSF Center for Advanced Light Microscopy (K. Herrington, S. Kim, and D. Larsen), the UCSF Center for Advanced Technology (E. Chow), and the UCSF Helen Diller Family Comprehensive Cancer Center Laboratory for Cell Analysis (S. Elmes; supported by the National Institutes of Health under award P30CA082103). These studies were supported by grants from the National Institutes of Health National Cancer Institute (F32CA260118 to E.E.B.), National Institute on Drug Abuse (DA010711 and DA012864 to M.v.Z.) and National Institute of Mental Health (MH120212 to M.v.Z.).

## Author Contributions

E.E.B. and M.v.Z. conceived of and designed the research. E.E.B. carried out experiments. E.E.B. and M.v.Z. analyzed data and wrote the paper.

## Competing Interests

The authors declare no competing interests.

## Data Availability

The data that support the findings of this study are available from the authors upon reasonable request. Source data and uncropped blots for Figs. 1–5, Extended Data Figs. 1–2 and 4-8 are provided with this paper. All plasmids and cell lines generated in this study are available from the authors.

## Notes

### Competing Interest Statement

The authors have declared no competing interest.

### Summary of Updates

This version of the manuscript has been revised to include new data and analysis.

