## Extended Data for "β-arrestin-independent endosomal cAMP signaling by a polypeptide hormone GPCR"

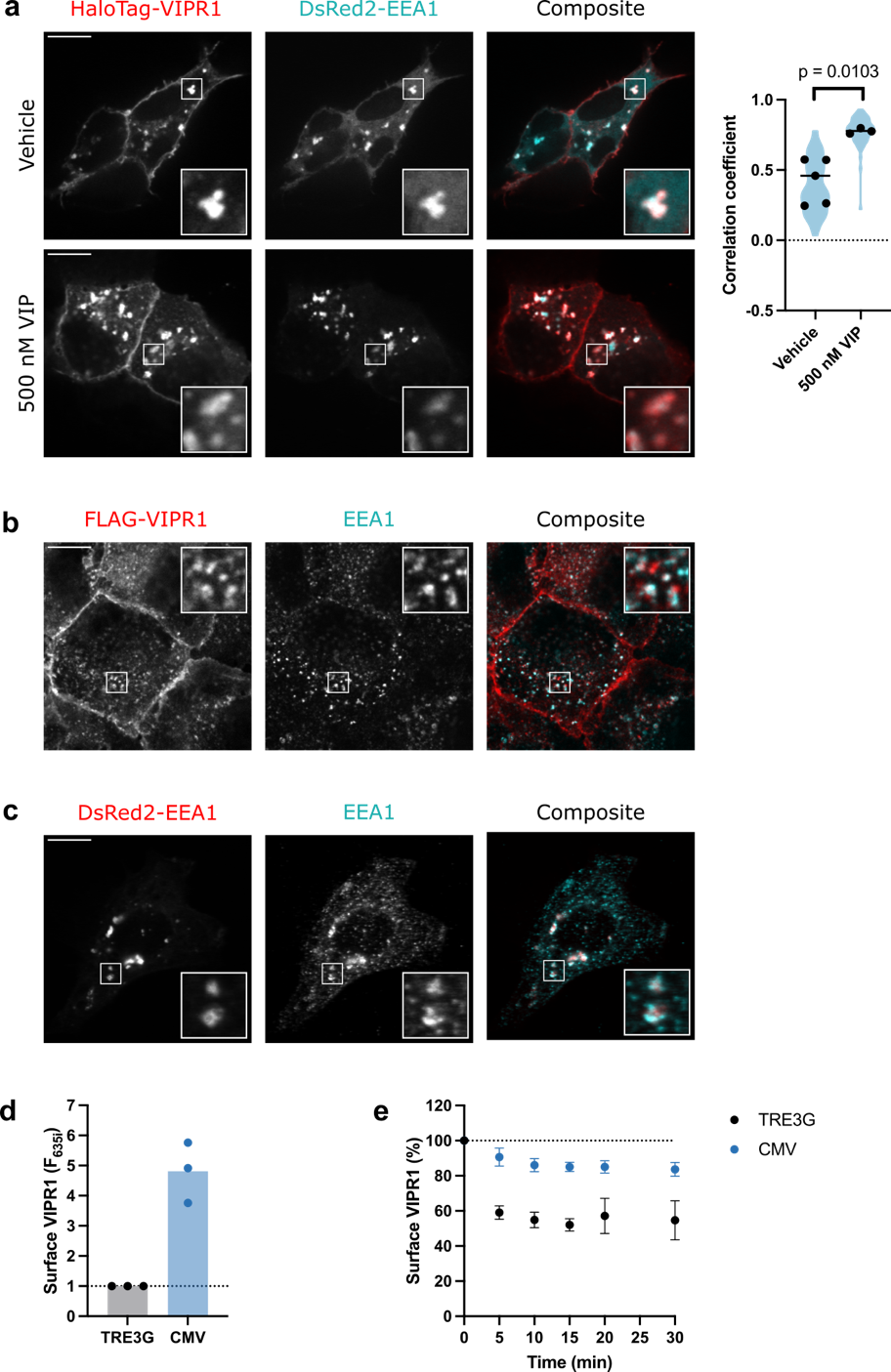

**Extended Data Fig. 1 VIPR1 undergoes agonist-induced internalization.** **a**, Cells coexpressing HaloTag-VIPR1, labeled with cell-impermeant JF_635_i-HTL for 10 minutes, and DsRed2-EEA1 were treated with vehicle or 500 nM VIP for 15 minutes before fixation and imaging. Scale bar is 10 μm. For quantification, cells were manually segmented, and the correlation coefficient was calculated for each cell. Mean correlation coefficients are shown for each biological replicate (9-27 cells), with a line at the mean of means and cell-level data superimposed as a violin plot. Significance was determined by an unpaired t-test. **b**, Representative image showing colocalization of FLAG-VIPR1 and endogenous EEA1 after a 15 minute treatment with 500 nM VIP prior to fixation. Surface FLAG-VIPR1 was labeled with anti-FLAG M1 antibody conjugated to Alexa Fluor 647 for 10 minutes before VIP treatment. Scale bar is 10 μm. **c**, Representative image showing colocalization of overexpressed DsRed-EEA1 and endogenous EEA1. Scale bar is 10 μm. **d**, Comparison of surface HaloTag-VIPR1 levels in cells stably expressing receptor under an inducible (TRE3G, black) or constitutive (CMV, blue) promoter, as measured by flow cytometry. **e**, Time course of surface HaloTag-VIPR1 expressed under an inducible (TRE3G, black) or constitutive (CMV, blue) promoter after treatment with 500 nM VIP, as measured by flow cytometry. Data for TRE3G is repeated from **Fig. 1b**. n = 3. For all panels, data represent biological replicates and are shown as individual data points or mean ± s.d.

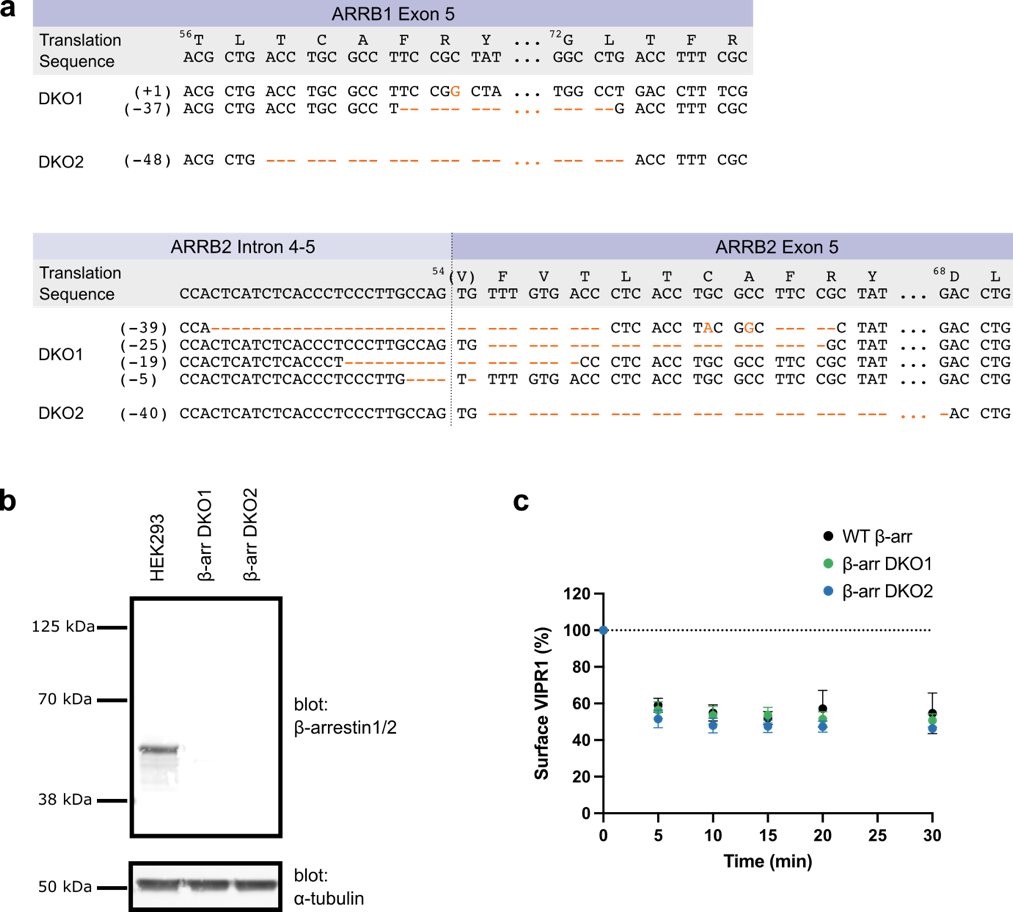

**Extended Data Fig. 2 Characterization of β-arr1/2 DKO CRISPR knockout cell lines used in this study.** **a**, Alignments showing the genetic modifications of β-arr1 (ARRB1) and β-arr2 (ARRB2) in independent monoclonal β-arrestin1/2 double knockout (β-arr1/2 DKO) cell lines. β-arr DKO2 is used for main text figures, while β-arr DKO1 is used for extended data figures. **b**, Western blot of parental HEK293 and β-arr1/2 DKO cell lysate, probing for β-arrestin1/2 and ɑ-tubulin as a loading control. **c**, Flow cytometry showing the time course of internalization of HaloTag-VIPR1 in WT (black), β-arr DKO1 (green), β-arr DKO2 (blue) and cells treated with 500 nM VIP. Data for WT β-arr is repeated from **Fig. 1b**. Data represent three biological replicates shown as mean ± s.d.

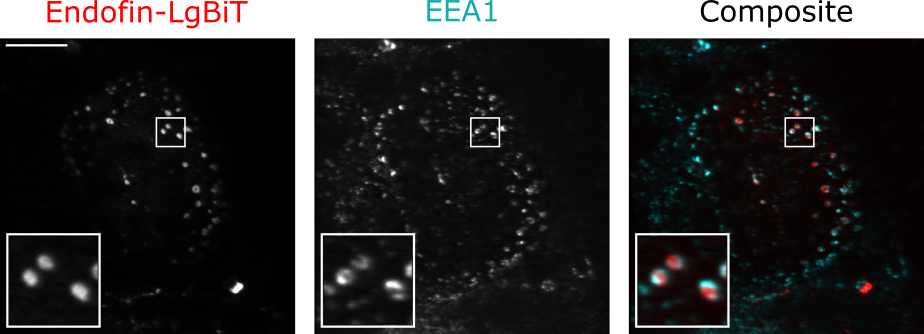

**Extended Data Fig. 3 Endofin-LgBiT and EEA1 localize to an overlapping population of endosomes.** Representative image showing overexpressed endofin-LgBiT, stained with an anti-LgBiT antibody, and endogenous EEA1. Scale bar is 10 μm.

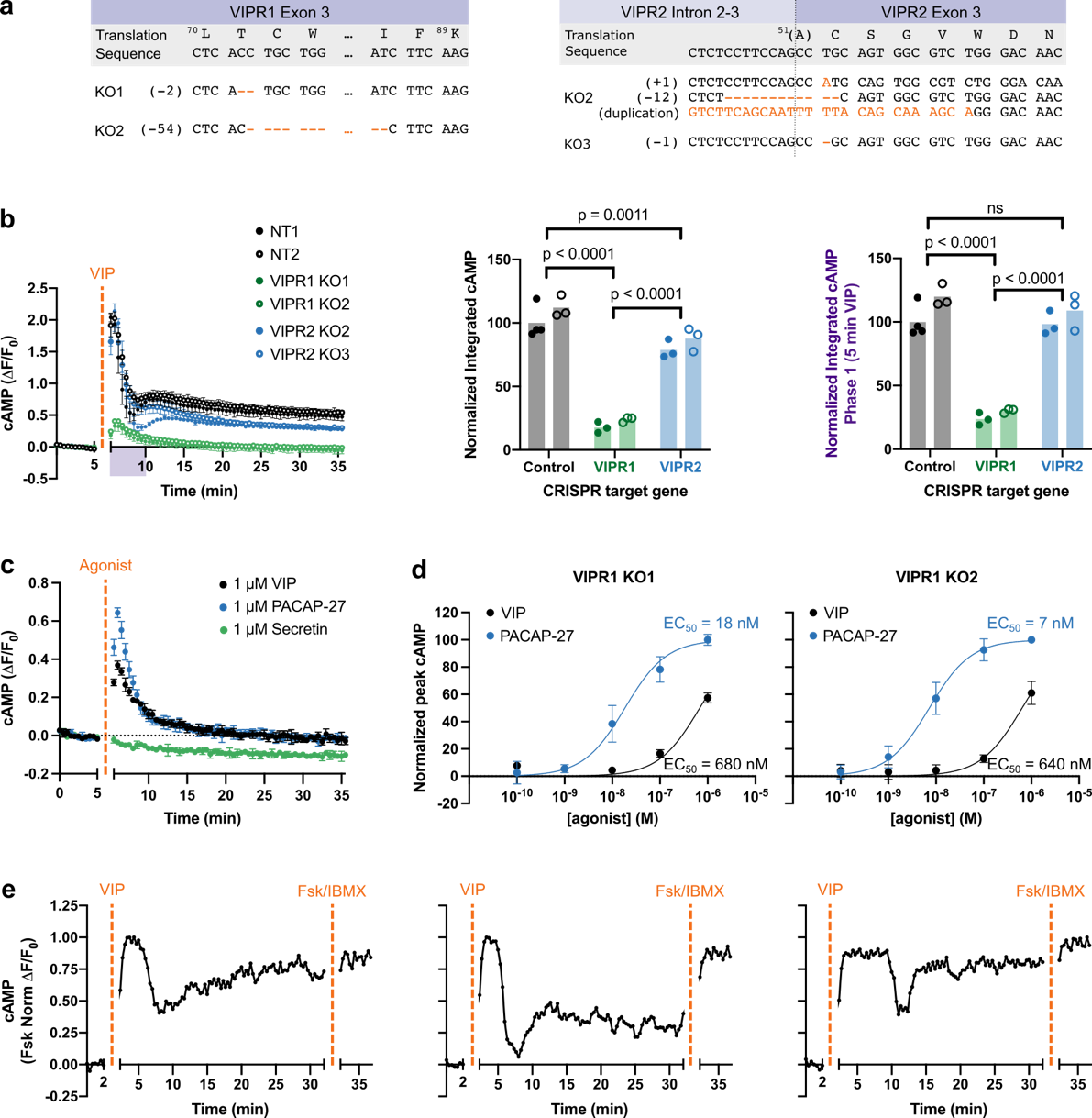

**Extended Data Fig. 4** **VIPR1 mediates both phases of cAMP signaling.** **a**, Alignments showing the genetic modifications of VIPR1 and VIPR2 in independent monoclonal lines. **b**, Changes in cAMP in control (NT, black), VIPR1 KO (green), and VIPR2 KO (blue) cell lines upon treatment with 500 nM VIP added at 5 minutes. Integrated cAMP of the entire timecourse and 0-5 min VIP treatment were calculated as the area under the curve and normalized to the average of NT1. Significance was determined by two-way ANOVAs with Tukey’s multiple comparisons tests. Closed and open circles refer to two independent clonal cell lines. **c**, Change in cAMP in VIPR1 KO1 cell line when stimulated with 1 µM VIP (black), PACAP (blue), or secretin (green). **d**, Dose response curves for stimulation of VIPR1 KO1 and KO2 cell lines with VIP (black) or PACAP (blue). Peak cAMP, normalized to the average maximum response elicited by 1 µM PACAP, is plotted. EC_50_ were calculated as noted in Methods. **e**, Example changes in cAMP in individual cells co-expressing mCherry upon treatment with 1 μM VIP added at 2 minutes and forskolin (Fsk, 10 µM) and IBMX (500 µM) at 32 minutes. Fluorescence was measured by microscopy and normalized to maximum fluorescence change of the ROI. Corresponding aggregate data is shown in **Fig. 2b**. For (**b-d)**, data represent three biological replicates shown as individual data points or mean ± s.d.

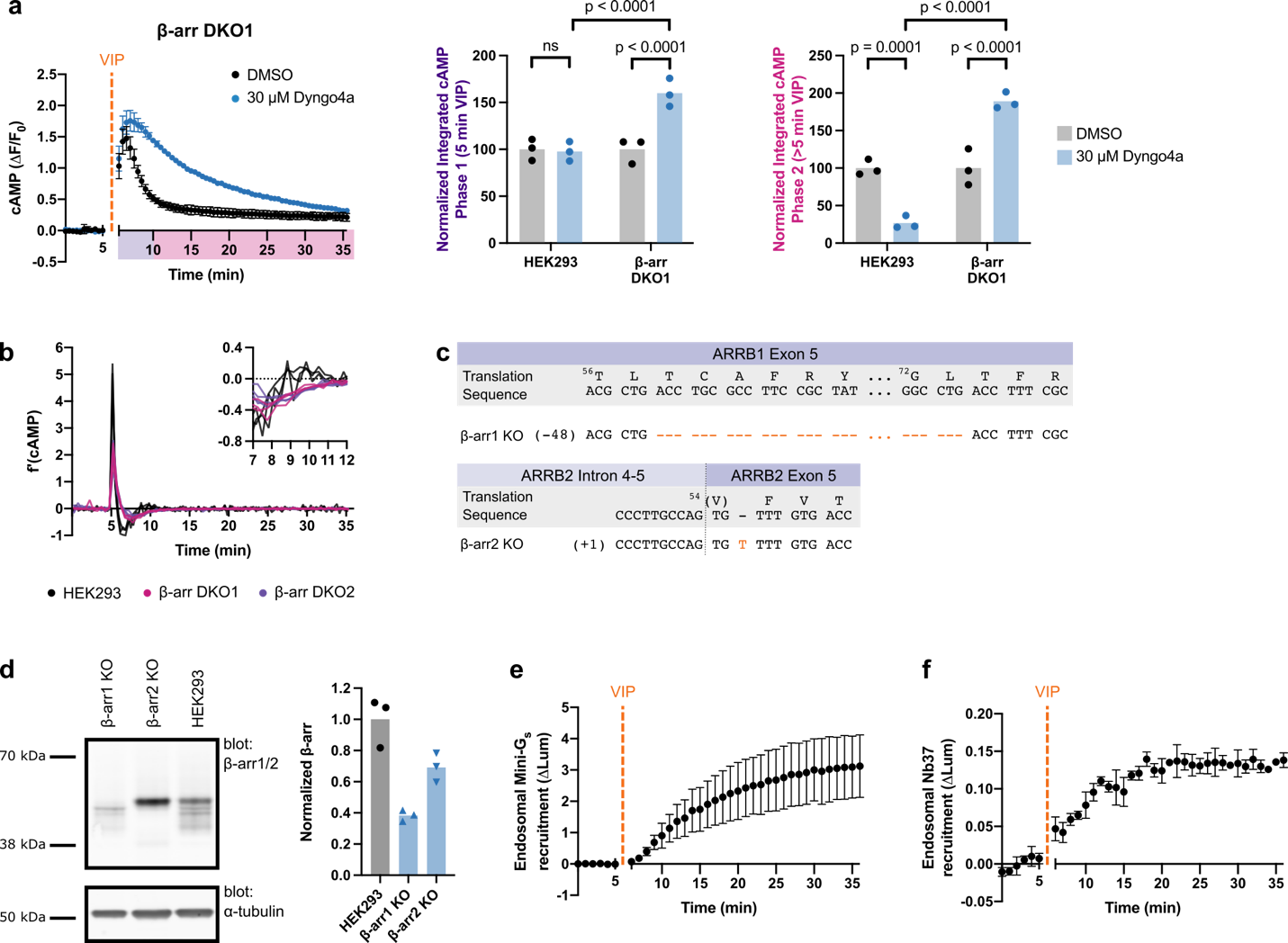

**Extended Data Fig. 5** **Independent β-arr1/2 DKO cell line recapitulates cAMP signaling trends.** **a**, Changes in cAMP in β-arr DKO1 cells upon treatment with 500 nM VIP added at 5 minutes. Integrated cAMP of each phase was calculated as the area under the curve and normalized to the average DMSO value for each cell line. Data for WT HEK293 repeated from **Fig. 2a**. Significance was determined by two-way ANOVA with Tukey’s multiple comparisons test along with data shown in **Fig. 3b**. n = 3. **b**, First derivatives of each cAMP replicate time course measured in HEK293 (black), β-arr DKO1 (pink), and β-arr DKO2 (purple) cells, with timepoints between 7-12 minutes highlighted in the right panel. Data from (**a**), **Fig. 2a**, and **Fig. 3a**. **c**, Alignments showing the genetic modifications of β-arr1 (ARRB1) and β-arr2 (ARRB2) in monoclonal knockout lines. **d**, Western blot of parental HEK293 and β-arr1/2 KO cell lysates, probing for β-arrestin1/2 and ɑ-tubulin as a loading control. Quantification of blots from three independent samples, normalizing to loading and the HEK293 parental cell line, is shown on right. **e-f**, NanoBiT bystander assays showing endosomal recruitment of mini-G_s_ (**e**, n = 4) or Nb37 (**f**, n = 3) upon addition of 1 µM VIP at 5 minutes in β-arr DKO1 cells. For all panels, data represent biological replicates and are shown as individual data points or mean ± s.d.

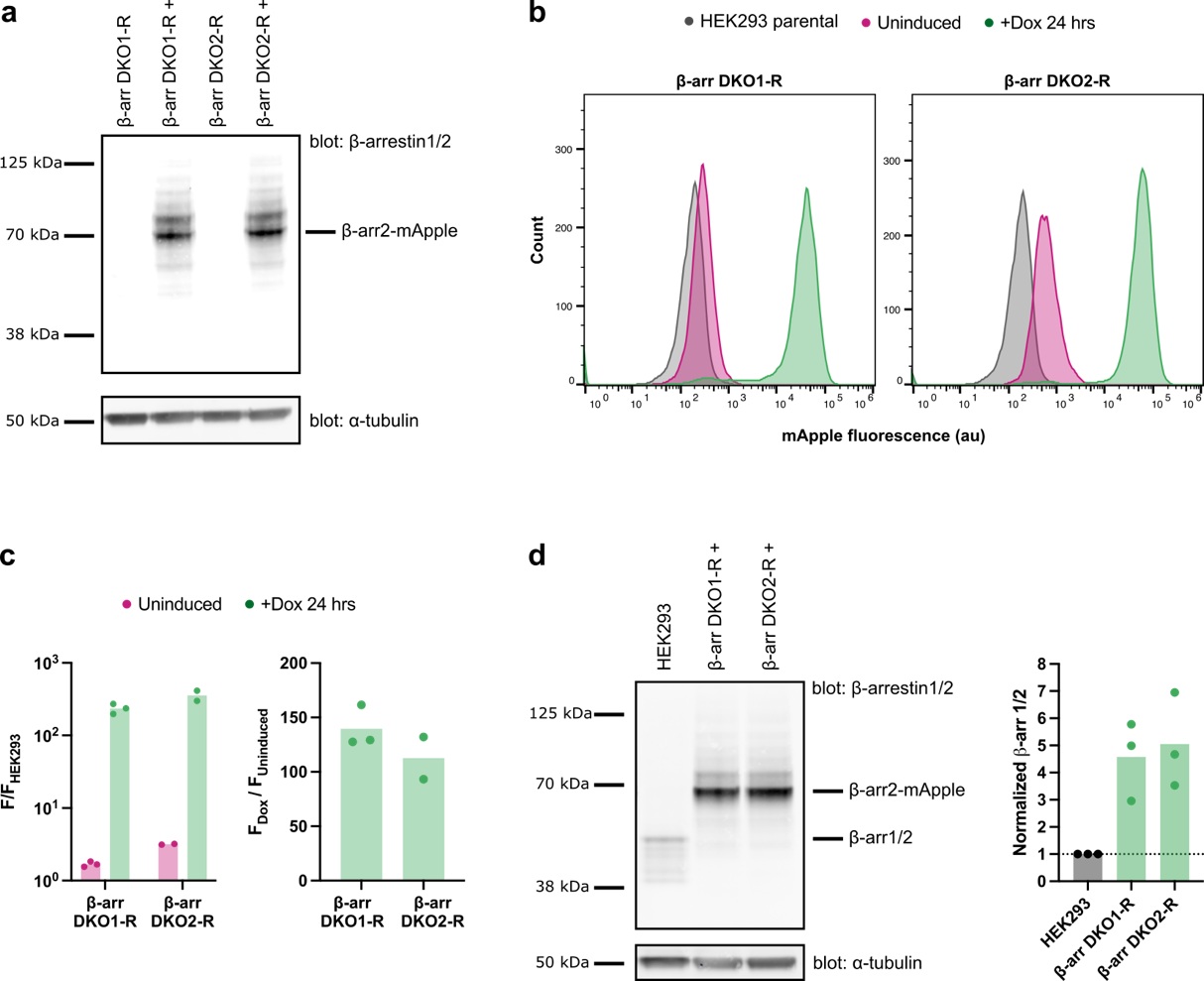

**Extended Data Fig. 6 Characterization of β-arr1/2 DKO rescue cell lines.** **a**, Western blot of β-arr1/2 DKO-R cell lysates, probing for β-arr1/2 and ɑ-tubulin as a loading control. Rescue cell lines stably express β-arr2-mApple under the control of a tet-inducible promoter (P_TRE3GS_). For induction (“+”), cells were treated with doxycycline (Dox) for 24 hours. **b**, Representative flow cytometry data showing the expression of β-arr2 upon dox induction, as measured by mApple fluorescence. **c**, Quantification of flow cytometry data shown in (**b**). Ratios of the median mApple fluorescence are shown compared to parental HEK293 cells (right) and the uninduced β-arr1/2 DKO-R cell lines (left). **d**, Western blot of parental HEK293 cells and induced β-arr1/2 DKO-R cells, probing for β-arr1/2 and ɑ-tubulin as a loading control. Quantification of blots from three independent inductions, normalizing to loading and the HEK293 parental cell line, is shown on right. For all panels, data represent biological replicates and are shown as individual data points.

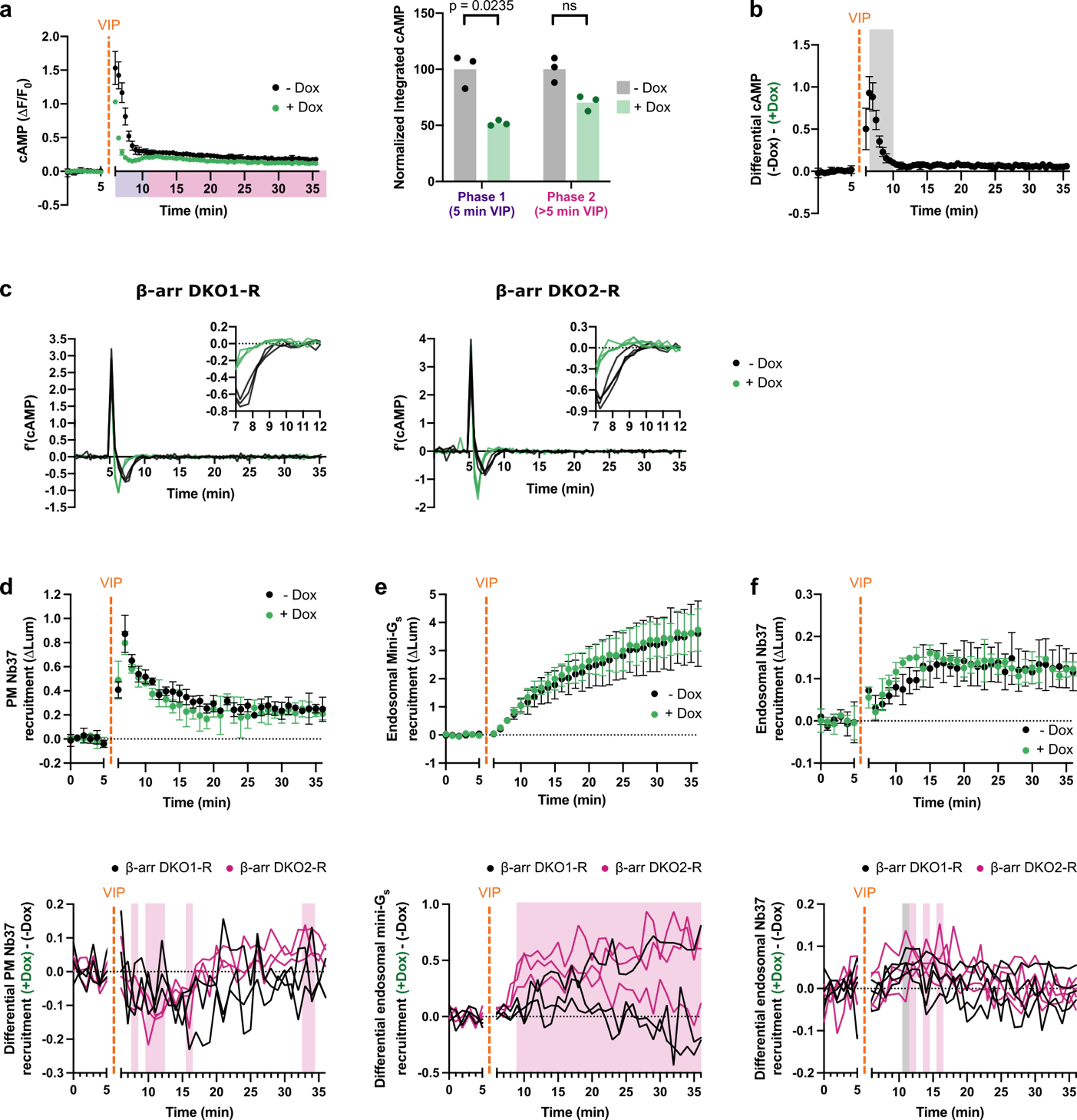

**Extended Data Fig. 7 Two independent β-arr1/2 DKO-R cell lines demonstrate β-arrestin’s role in desensitizing the first phase of VIPR1 signaling. a**, Changes in cAMP in β-arr DKO1-R cells with (green) and without (black) Dox induction upon treatment with 500 nM VIP added at 5 minutes. Integrated cAMP of each phase (0-5 min or 5-30 min VIP treatment) was calculated as the area under the curve and normalized to the average -Dox value. Significance was determined by a repeated measures 2 way ANOVA with Sidak's multiple comparisons test along with data for DKO2-R shown in **Fig. 4a.** n = 3. **b**, Difference in cAMP response with and without β-arr2 expression, calculated as the difference in fluorescence change (ΔF/F_0_) between the -Dox and +Dox curves shown in (**a**). Shaded area represents timepoints where the difference is statistically significant, as determined by a repeated measures 2 way ANOVA with Sidak's multiple comparisons test (p < 0.05) carried out on the curves in (**a**). **c**, First derivatives of each cAMP replicate time course measured in β-arr DKO2-R (**Fig. 4a**) and β-arr DKO1-R (**a**). **d-f**, NanoBiT bystander assays showing plasma membrane recruitment of Nb37(**d**, n = 3) or endosomal recruitment of mini-G_s_ (**e**, n = 3) and SmBiT (**f**, n = 4) in β-arr DKO1-R cells upon addition of 1 µM VIP at 5 minutes. Shown below each panel is the differential recruitment, calculated as the difference between +Dox and -Dox curves, for both β-arr DKO1-R (black) and β-arr DKO2-R (pink, **Fig. 4c-e**) cell lines. Shaded areas represent timepoints where the difference is statistically significant, as determined by a repeated measures 2 way ANOVA with Sidak's multiple comparisons test (p < 0.05), for β-arr DKO1-R (black) and β-arr DKO2-R (pink). For all panels, cells treated with Dox to induce expression of β-arr2-mApple are shown in green, and data represent biological replicates, shown as individual data points or mean ± s.d.

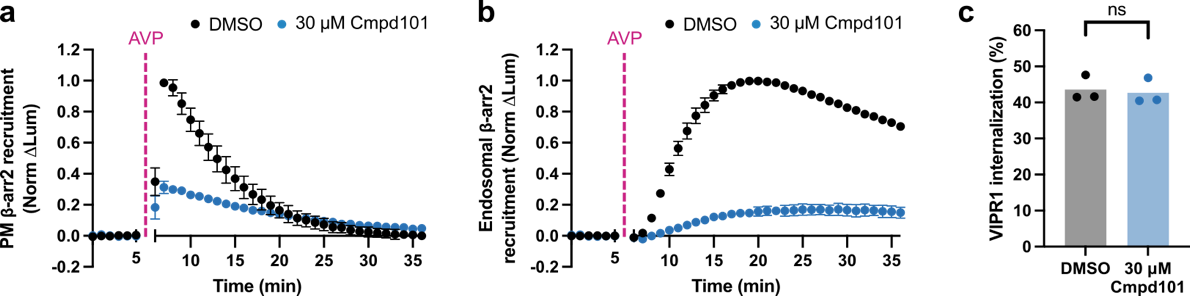

**Extended Data Fig. 8 Cmpd101 differentially modulates receptor-β-arrestin complexes. a-b**, NanoBiT bystander assays showing plasma membrane (**a**) or endosomal (**b**) recruitment of β-arr2 in HEK293 cells overexpressing V2R upon addition of 1 µM AVP at 5 minutes. **c**, Internalization of HaloTag-VIPR1 after a 30 minute treatment with 500 nM VIP, as measured by flow cytometry. Significance was determined by a paired t-test.

**Extended Data Tables**

**Extended Data Table 1 Estimated decay rates for the first phase of endogenous VIPR1-mediated cAMP signaling.**

| **Cell Line** | **Treatment** | **k (s^-1^)** | **95% CI (s^-1^)** |
| --- | --- | --- | --- |
| HEK293 | DMSO | 0.63 | 0.26 - 1.04 |
|  | 30 µM Dyngo4a | 0.49 | 0.45 - 0.53 |
| β-arr DKO1 | DMSO | 0.45 | 0.38 - 0.53 |
|  | 30 µM Dyngo4a | 0.097 | 0.093 - 0.10 |
| β-arr DKO2 | DMSO | 0.38 | 0.26 - 0.52 |
|  | 30 µM Dyngo4a | 0.019 | undetermined |
| β-arr1 KO | DMSO | 1.36 | 0.89 - 1.94 |
|  | 30 µM Dyngo4a | 0.83 | 0.72 - 0.95 |
| β-arr2 KO | DMSO | 0.72 | 0.49 - 0.98 |
|  | 30 µM Dyngo4a | 0.52 | 0.47 - 0.58 |
| β-arr DKO1-R | - | 0.90 | 0.81 - 0.99 |
|  | Dox (24 hr) | 1.85 | 1.67 - 2.06 |
| β-arr DKO2-R | - | 1.29 | 0.94 - 1.80 |
|  | Dox (24 hr) | 2.24 | 1.92 - 2.61 |
