## Supplemental Data for "β-arrestin-independent endosomal cAMP signaling by a polypeptide hormone GPCR"

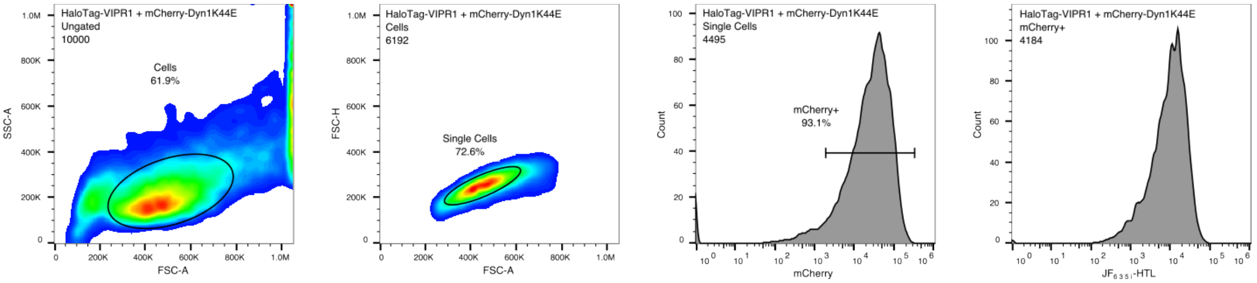


**Supplementary Fig. 1 Example flow cytometry gating strategy.** Gating strategy for cells co-expressing HaloTag-VIPR1 and mCherry/mCherry-Dyn1K44E is shown. For all other experiments with cells expressing only a HaloTag receptor, the gating strategy is the same, excluding the mCherry+ gate.

**Supplementary Table 1 DNA constructs used in this study.**

| Construct | Source |
|---|---|
| pcDNA3_mCherry | This study: deletion of Dyn1 from pcDNA_mCherry-Dyn1[1] |
| pcDNA3_mCherry-Dyn1K44E | [1] |
| pCMV-Dest_mCherry | This study; insertion of mCherry into pCMV-Dest (Invitrogen A24223) |
| pCMV-Dest_mCherry-Dyn1K44E | This study; insertion of mCherry-Dyn1K44E[1] into pCMV-Dest (Invitrogen A24223) |
| pCAGGS_LgBiT-CAAX | [2] |
| pCAGGS_endofin-LgBiT | [2] |
| pcDNA3.1_SS-HaloTag-VIPR1-LgBiT | This study: C-terminal insertion of LgBiT into pcDNA3.1_SS-HaloTag-VIPR1. |
| pEGFP-βarr2-SmBiT | [3] |
| pcDNA3.1_SmBiT-miniGs | This study: insertion of miniGs[4] into pcDNA3.1 with SmBiT(114) added into primers. |
| pcDNA3.1_Nb37-SmBiT(114) | This study: insertion of Nb37 (from R. Irannejad) into pcDNA3.1 with SmBiT(114) added into primers. |
| pcDNA3.1_Nb37-SmBiT(101) | This study: insertion of Nb37 (from R. Irannejad) into pcDNA3.1 with SmBiT(101)[5] added into primers. |
| GNB1-T2A-cpVenus-GNGT1-IRES-GNAS | This study: deletion of Nluc from Gs(short)-CASE. Gs(short)-CASE was a gift from Gunnar Schulte (Addgene plasmid # 168124). |
| pcDNA3.1_SS-FLAG-VIPR1 | This study: insertion of VIPR1(31-457) (from A. Marley) into pcDNA3.1 following the signal sequence of influenza hemagglutinin (SS) and FLAG-linker (MKTIIALSYIFCLVFADYKDDDDGGS) |
| pcDNA3.1_SS-HaloTag-VIPR1 | This study: replacement of FLAG-linker with HaloTag7 in pcDNA3.1-SS-HaloTag-VIPR1 |
| pcDNA_SS-HaloTag-β2AR | This study: replacement of FLAG with HaloTag7 in pcDNA_SS-FLAG-β2AR[6] |
| pcDNA_SS-FLAG-V2R | From A. Lazar |
| pDsRed2-EEA1(2xFYVE) | From R. Irannejad |
| pmApple_Arr2-mApple | [7] |
| pLVX-TetOne-Puro_Arr3-mApple | This study; insertion of Arr3-mApple[7] into pLVX-TetOne-Puro (Takara Bio). |

**Supplementary Table 2 CRISPR knockout cell line reagents.** Two sets of validation primers reflect a nested PCR strategy.

| Target | Cell line(s) | sgRNA sequence | Forward PCR validation primer(s) | Reverse PCR validation primer(s) |
|---|---|---|---|---|
| VIPR1 | VIPR1 KO1 & KO2 | UGUGGGACAACCUCACCUGC | GTGAGCCTTTCCCACATGCTTATC | CAACAGTCAAAACCACCACTGGC |
| VIPR2 | VIPR2 KO1 & KO2 | UCCCAGACGCCACUGCAGGC | GAAGGCAGGTGATGCCCTTCT | CCAGTGGTGCCCATTTCATCACA |
| ARRB1 | β-arr1 KO, β-arr DKO1 & DKO2 | GUCCUCCCGGCCAUAGCGGA | CATACCCCTCCACATATGCCC | GTCCCAGCATACACTGGGAAAC |
| ARRB2 | β-arr2 KO, β-arr DKO1 & DKO2 | CAGGUGAGGGUCACAAACAC | 1. GGATGAGAAGGGAAGAAGGAAGG<br>2.CTGCCTCACTGTTTCTTCCAGATC | 1.GACAAAGTGGACCCTGTAGGTAAG<br>2.CAGGAAGTGAGCTGGTGTGTC |
| ARRB2 | β-arr DKO2 | CCGCUAUGGCCGUGAAGACC | 1. GGATGAGAAGGGAAGAAGGAAGG<br>2. CTGCCTCACTGTTTCTTCCAGATC | 1. GACAAAGTGGACCCTGTAGGTAAG<br>2. CAGGAAGTGAGCTGGTGTGTC |
| NT1 | NA | GCACUACCAGAGCUAA | NA | NA |
| NT2 | NA | GUACGUCGGUAUAACU | NA | NA |

### Supplementary Table 3 CRISPR NGS sequencing primers

| Target | Forward primer | Reverse primer |
|---|---|---|
| ARRB2 | ACACTCTTTCCCTACACGACGCTCTTCC GATCTAAGTGAAATGGGCTGGCCTTGG | GACTGGAGTTCAGACGTGTGCTCTTC CGATCTATGCTGGCCCAGCTTCCTCA |

**Supplementary Table 4. List of antibodies used.**

| Antibody | Manufacturer | Catalog # | Use | Dilution |
|---|---|---|---|---|
| Rabbit anti β-arrestin 1/2 (D24H9) | Cell Signaling Technologies | 4674 | WB | 1:1000 |
| Mouse anti α-tubulin | Cell Signaling Technologies | 3873 | WB | 1:1000 |
| IRDye 680RD Donkey anti-Mouse IgG | LI-COR Biosciences | 926-68072 | WB | 1:10000 |
| IRDye 800CW Donkey anti-Rabbit IgG | LI-COR Biosciences | 926-32213 | WB | 1:10000 |
| Mouse anti LgBiT | Promega | N7100 | IF | 1:250 |
| Goat anti EEA1 (N-19) | Santa Cruz Biotechnology | sc-6415 | IF | 1:250 |
| Donkey anti goat IgG AlexaFluor 488 | Invitrogen | A-11055 | IF | 1:500 |
| Donkey anti mouse IgG AlexaFluor 647 | Invitrogen | A-31571 | IF | 1:500 |
